# Chemical genomics informs antibiotic and essential gene function in *Acinetobacter baumannii*

**DOI:** 10.1101/2024.12.05.627103

**Authors:** Jennifer S Tran, Ryan D Ward, Rubén Iruegas-López, Ingo Ebersberger, Jason M Peters

## Abstract

The Gram-negative pathogen, *Acinetobacter baumannii*, poses a serious threat to human health due to its role in nosocomial infections that are resistant to treatment with current antibiotics. Despite this, our understanding of fundamental *A. baumannii* biology remains limited, as many essential genes have not been experimentally characterized. These essential genes are critical for bacterial survival and, thus, represent promising targets for drug discovery. Here, we systematically probe the function of essential genes by screening a CRISPR interference knockdown library against a diverse panel of chemical inhibitors, including antibiotics. We find that most essential genes show chemical-gene interactions, allowing insights into both inhibitor and gene function. For instance, knockdown of lipooligosaccharide (LOS) transport genes increased sensitivity to a broad range of chemicals. Cells with defective LOS transport showed cell envelope hyper-permeability that was dependent on continued LOS synthesis. Using phenotypes across our chemical-gene interaction dataset, we constructed an essential gene network linking poorly understood genes to well-characterized genes in cell division and other processes. Finally, our phenotype-structure analysis identified structurally related antibiotics with distinct cellular impacts and suggested potential targets for underexplored inhibitors. This study advances our understanding of essential gene and inhibitor function, providing a valuable resource for mechanistic studies, therapeutic strategies, and future key targets for antibiotic development.

## Introduction

Systems biology provides a robust framework for deciphering the complex networks that govern cellular functions. This approach is especially pertinent in the study of infectious diseases and antibiotic resistance as it offers sophisticated tools to analyze the multifaceted interactions between pathogens and their environments, including their responses to therapeutic interventions. The Gram-negative, hospital-acquired pathogen *Acinetobacter baumannii* is categorized as an ‘urgent threat’ due to certain clinical strains having developed resistance to all known therapeutics and its ability to persist on surfaces that are typically adverse to cellular life, such as stainless steel (1, 2). Despite the critical dangers posed by *A. baumannii*, our understanding of how the fundamental elements of its biology interact with antibiotics or other inhibitors remains limited. Essential genes, which are vital for survival even under optimal growth conditions, are promising targets for drug discovery. Essential genes in *A. baumannii* have been cataloged using methods like transposon sequencing (Tn-seq), which identifies genes with low or nonexistent insertion frequencies as critical for survival (3, 4). However, lethality caused by knocking out essential genes limits more detailed studies of gene function and interactions. CRISPR interference (CRISPRi), which allows for the knockdown of gene expression without eliminating gene function (5, 6), provides a solution for studying the function of essential genes (7). CRISPRi uses a deactivated Cas9 protein, dCas9, directed by a single guide RNA (sgRNA) to specifically target genes for knockdown. This method has been used successfully in diverse bacteria including *A. baumannii*, non-*baumannii Acinetobacter* species, and other Gram-negative pathogens (8–13).

Chemical genomics, which combines gene perturbation libraries with chemical treatments, offers a strategy to explore gene function as changes in relative fitness of mutant or knockdown cells in response to chemical stresses (14, 15). This approach has been applied at the genome-scale in bacteria, facilitating the observation of subtle phenotypic responses and the identification of genetic connections from Tn-seq or deletion libraries (3, 16, 17) as well as in CRISPRi libraries (18–21). Our previous work involved developing and characterizing a CRISPRi library in *A. baumannii*, which identified key vulnerabilities among essential genes and demonstrated how this library could be utilized to elucidate antibiotic-gene interactions (20).

In this study, we took a chemical-genomics approach that provided systems-level insights into gene and antibiotic function in *A. baumannii*. We screened our CRISPRi library against a set of 45 chemical stressors, allowing us to identify pathways that cause broad sensitivity to these agents when perturbed. Additionally, we created an essential gene network that informed the function of poorly characterized genes that are unique to or highly divergent in *A. baumannii*.

Finally, we integrated phenotypic and chemoinformatic datasets to identify possible target pathways for inhibitors and show distinctions in the physiological impacts of structurally similar drugs. In doing so, this work advances our understanding of the cellular contributions of *A. baumannii* essential genes in the context of antibiotic stress.

## Results

### The vast majority of A. baumannii essential genes show significant chemical-gene interactions

To identify phenotypes for essential gene knockdown strains in the context of chemical treatment (i.e., chemical-gene interactions) in *A. baumannii*, we conducted a large-scale chemical genomics screen using our established inducible CRISPRi library in *A. baumannii* strain ATCC19606 (19606 throughout) (20). This library consists of pooled CRISPRi strains with guides targeting 406 putatively essential genes derived from Tn-seq data in other strains (3, 4), along with 1000 non-targeting control sgRNAs. For each targeted gene, we cloned four sgRNA spacers that exactly match the target sequence (perfect-match) and ten mismatch spacers with single-base variations from the target sequence that allow for titration of knockdown (22). We performed competition fitness assays by inducing CRISPRi knockdown and adding chemicals at sublethal concentrations. These chemical concentrations ensured sufficient cell viability to preserve library diversity and representation, allowing us to measure phenotypic responses to treatment (Fig 1A). We screened the library against a diverse collection of chemical compounds, including clinically relevant antibiotics, heavy metals, and compounds with unknown mechanisms of action (Fig 1B). Knockdown strain abundance was analyzed by amplification and sequencing of the sgRNA spacer regions. We improved quantification of sgRNA spacers by implementing a position-aware counter that ensures count accuracy for guides with heavily overlapping targets. We also set quality control metrics—replicate correlation and sample complexity cutoffs—to remove samples with excessive noise or selective pressure (e.g., lethal chemical concentrations) that would skew calculations (Appendix Fig S1A and B; see Material and Methods for details). We verified that chemical solvent impacts were negligible as library composition for DMSO-only and mock treatment samples were highly correlated (*r* > 0.95) (Appendix Fig S1C).

**Figure 1.**
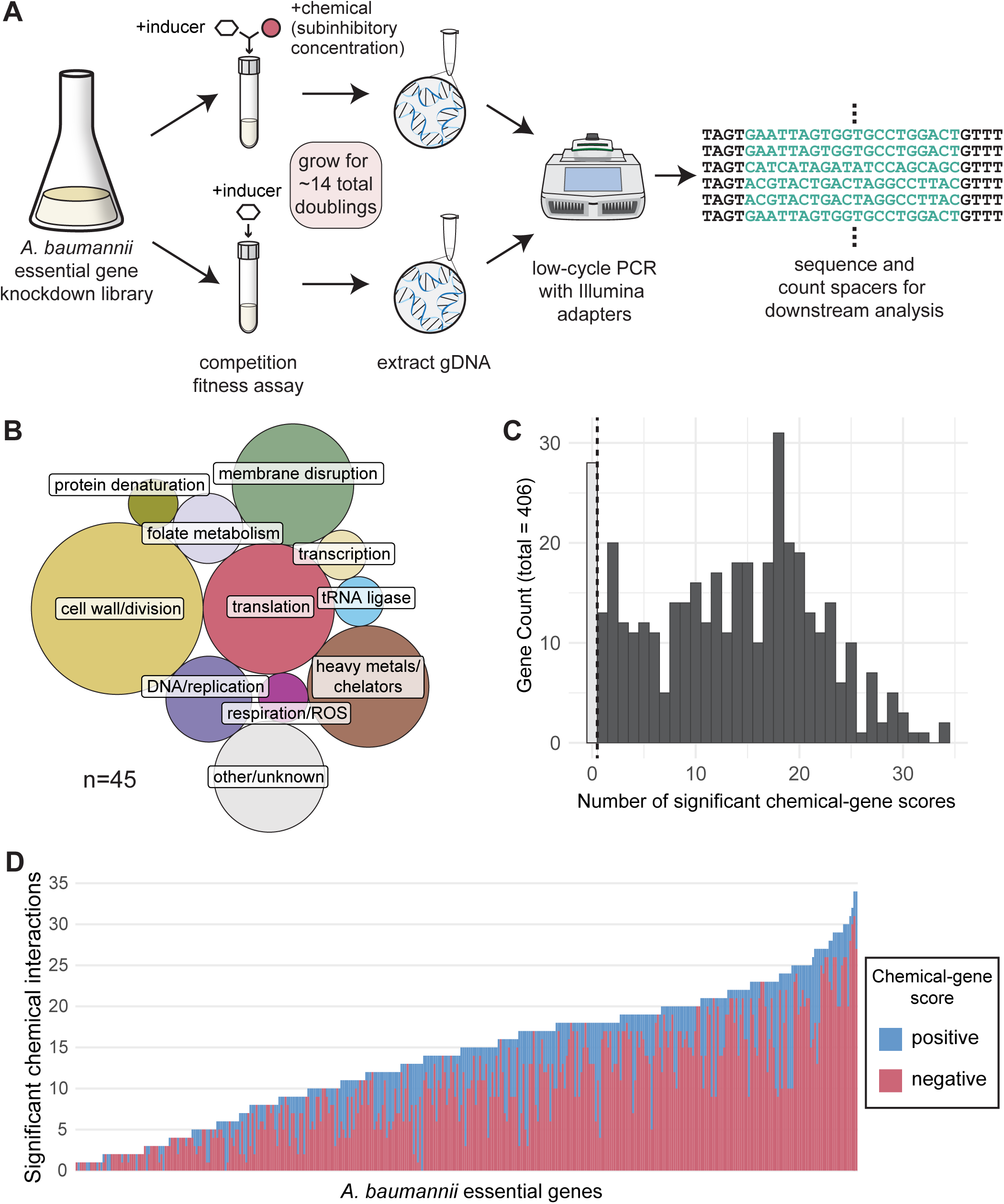
Chemical genomics screen in *A. baumannii* essential gene library. (A) Schematic depicting experimental setup for chemical genomics. (B) Proportional area chart of reported cellular targets of screen chemicals. (C) Histogram showing significant chemical-gene scores (|median log_2_ fold change| ≥ 1 and Stouffer’s *p* < 0.05, calculated from perfect-match guides), across chemical conditions in our screen. Darker bars to the right of the dotted line represent at least one significant chemical interaction in this screen. (D) Stacked bar chart showing the sum of significant positive (pink) and negative (blue) CG scores across screen chemicals for each gene in the library.

To determine chemical-gene phenotypes from the pooled screens, we calculated chemical-gene (CG) scores for each gene as the median log_2_ fold change (medL_2_FC) of perfect guides with chemical treatment compared to induction alone. Positive CG scores indicate gene knockdowns with improved growth under chemical conditions relative to the untreated knockdown, whereas negative CG scores indicate chemical sensitivity or reduced growth. 93% (378/406) of the genes we investigated exhibited at least one significant CG score (medL_2_FC ≥ |1|, *p* <0.05) upon knockdown, with a median of 14 significant chemical interactions per gene (Fig 1C). Most of these scores were skewed toward reduced growth, with ∼73% (3895/5345) of significant CG scores being negative (Fig 1D). This is consistent with previous studies suggesting essential genes are more frequently connected by negative interactions, where two perturbations (e.g., a chemical stressor and a knockdown) cause further loss of fitness than expected based the effects of the individual perturbations (23). Overall, this dataset provides thousands of unique chemical-gene phenotypes and serves as a detailed resource for *A. baumannii* essential gene responses to chemical stress.

### Lipooligosaccharide transport inhibition enhances drug susceptibility through increased membrane permeability

To identify *A. baumannii* pathways crucial for chemical resistance, we ranked genes based on significant negative CG scores (medL_2_FC ≤-1, *p* <0.05) across conditions and performed functional enrichment using the STRING database (24). We found that knockdowns of the lipooligosaccharide transport (Lpt) system were significantly enriched (GO:0015920, enrichment score = 8.95, FDR = 1.04e-05), with negative CG scores for 70% of conditions in our screen (Fig 2B and C). Lpt is a highly conserved pathway required for lipopolysaccharide (LPS)/lipooligosaccharide (LOS) export to the outer leaflet of the outer membrane in Gram-negative bacteria. *A. baumannii* uses Lpt to transport LOS, which is comprised of lipid A and core oligosaccharide moieties found in LPS but lacks O-polysaccharide repeats (Fig 2A) (25).

**Figure 2.**
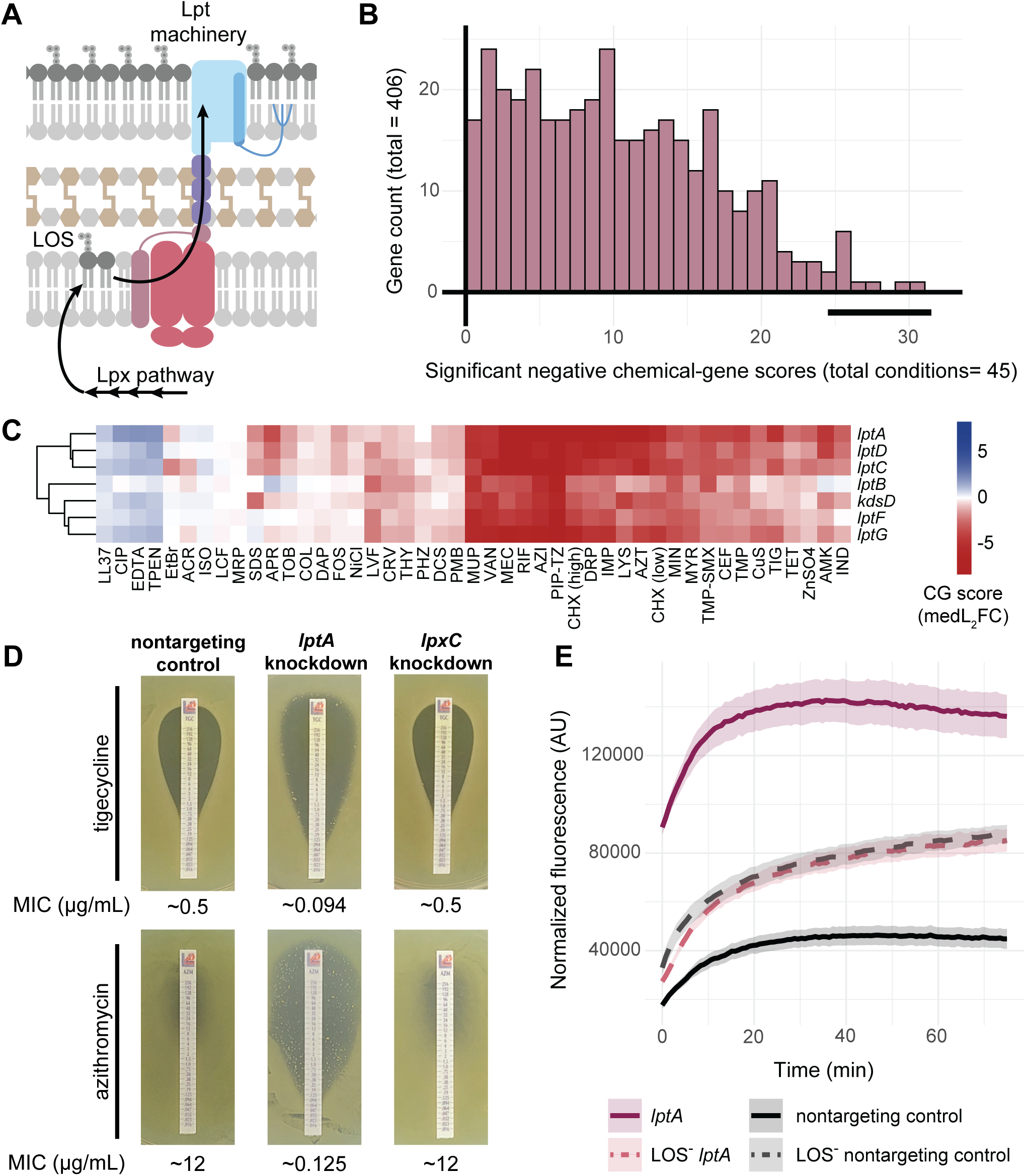
*lpt* knockdowns are sensitized to antibiotics. (A) Graphical depiction of Lpt machinery and LOS trafficking to the outer membrane in *A. baumannii*. (B) Histogram of significant negative chemical interactions (medL_2_FC ≤ −1 and Stouffer’s *p* < 0.05) across conditions. Bold line indicates location of *lpt* genes. (C) Heatmap displaying CG scores for *lpt* knockdown cluster. Y-axis clustering was conducted using the Ward method and Canberra distance across all library knockdowns. (D) Tigecycline and azithromycin MIC test strip assays for the *lptA* knockdown, *lpxC* knockdown, and control strain containing a nontargeting guide. Plates supplemented with 1mM IPTG; approximate MICs reported below images (μg/mL). (E) Ethidium bromide permeability assay for *lptA* knockdown and nontargeting control in 19606 or LOS^-^ backgrounds after induction; increased fluorescence over time indicates membrane permeability. Ribbons represent standard deviation (n=4).

Intriguingly, while LPS/LOS is essential in most Gram-negative bacteria, many strains of *A. baumannii* do not require LOS for survival, including the strain used for our library (19606) (26). Indeed, knockdowns of *lpx* genes responsible for LOS synthesis showed fewer phenotypes than knockdowns of *lpt* needed for LOS transport (Fig EV1A), suggesting that LOS transport unexpectedly plays a more crucial role in *A. baumannii*’s chemical sensitivity than LOS synthesis itself.

We validated our screen results for individual knockdowns outside of the pooled context by performing quantitative minimum inhibitory concentration (MIC) assays using antibiotic strips. This revealed that an *lptA* knockdown was sensitized to several clinically relevant antibiotics (Fig 2D). In contrast, an *lpxC* knockdown was not sensitized (Fig 2D and EV1B) although loss of LOS has been shown to increase antibiotic sensitivity (27). Both the *lpxC* and *lptA* knockdowns remained sensitized to colistin (Fig EV1B)—a cationic lipopeptide that disrupts membranes by targeting the lipid A moiety of LOS (28) —suggesting that these knockdowns do not completely eliminate LOS production or transport and knockdown strains still maintain some level of LOS in the outer membrane that can be targeted by colistin (29). We therefore introduced CRISPRi either targeting *lptA* or containing a non-targeting control sgRNA into a mutant strain background that does not produce LOS (i.e., LOS^-^), 19606 *lpxC(S106R*) (27). In contrast to the 19606 *lptA* and *lpxC* knockdowns, LOS^-^ strains were fully resistant to colistin (Fig EV1B).

Additionally, the LOS^-^ strains showed considerable growth defects not exhibited by the 19606 knockdown strains (Fig EV1D). Yet, the *lptA* knockdown strain was as similarly sensitized to levofloxacin as the LOS^-^ strains (Fig EV1B). This suggested levofloxacin, an antibiotic targeting gyrase/DNA synthesisin the cell cytoplasm, could penetrate the cell envelope more effectively in both the *lptA* and the LOS^-^ strains.

These findings along with the sensitivity of *lpt* knockdowns to chemically diverse inhibitors pointed to a permeability defect. Therefore, we performed an ethidium bromide (EtBr) fluorescence assay to test for the knockdown’s effect on membrane permeability; in this assay, cells are briefly exposed to EtBr, which fluoresces after passing through the outer and inner membranes and binding to DNA. More permeable membranes will have higher fluorescence readings, and as expected, the LOS^-^ strains showed increased permeability compared to the 19606 nontargeting control. Surprisingly though, the 19606 *lptA* knockdown was substantially more permeable than the LOS^-^ and LOS^-^ *lptA* knockdown strains (Fig 2E and EV1E). This indicates that Lpt deficiencies cause a further increase in membrane permeability when LOS is being synthesized, possibly due to LOS accumulating in and destabilizing the inner membrane. Taken together, these findings further underscore the critical role of the LOS transport system in *A. baumannii* in maintaining membrane integrity and mediating resistance to chemicals.

### Taxon-specific determinants of chemical susceptibility and resistance

Essential genes that are highly divergent in, or exclusive to, *A. baumannii* may contribute to its unique drug resistance and could help guide the development of targeted therapeutics. To identify these genes, we determined the phylogenetic profiles of the genes represented in our library across a broad collection species covering the known diversity of *Gammaproteobacteria* (30). We identified 15 candidate genes that were highly responsive to chemicals when knocked down but were rare in *Gammaproteobacteria* outside of the order to which *A. baumannii* belongs (Fig 3A; Appendix Fig S3). These genes exhibited significant CG scores (medL_2_FC ≥ |1|, *p* <0.05) for at least 10% of chemicals tested and likely play substantial roles in responding to chemical stressors.

**Figure 3.**
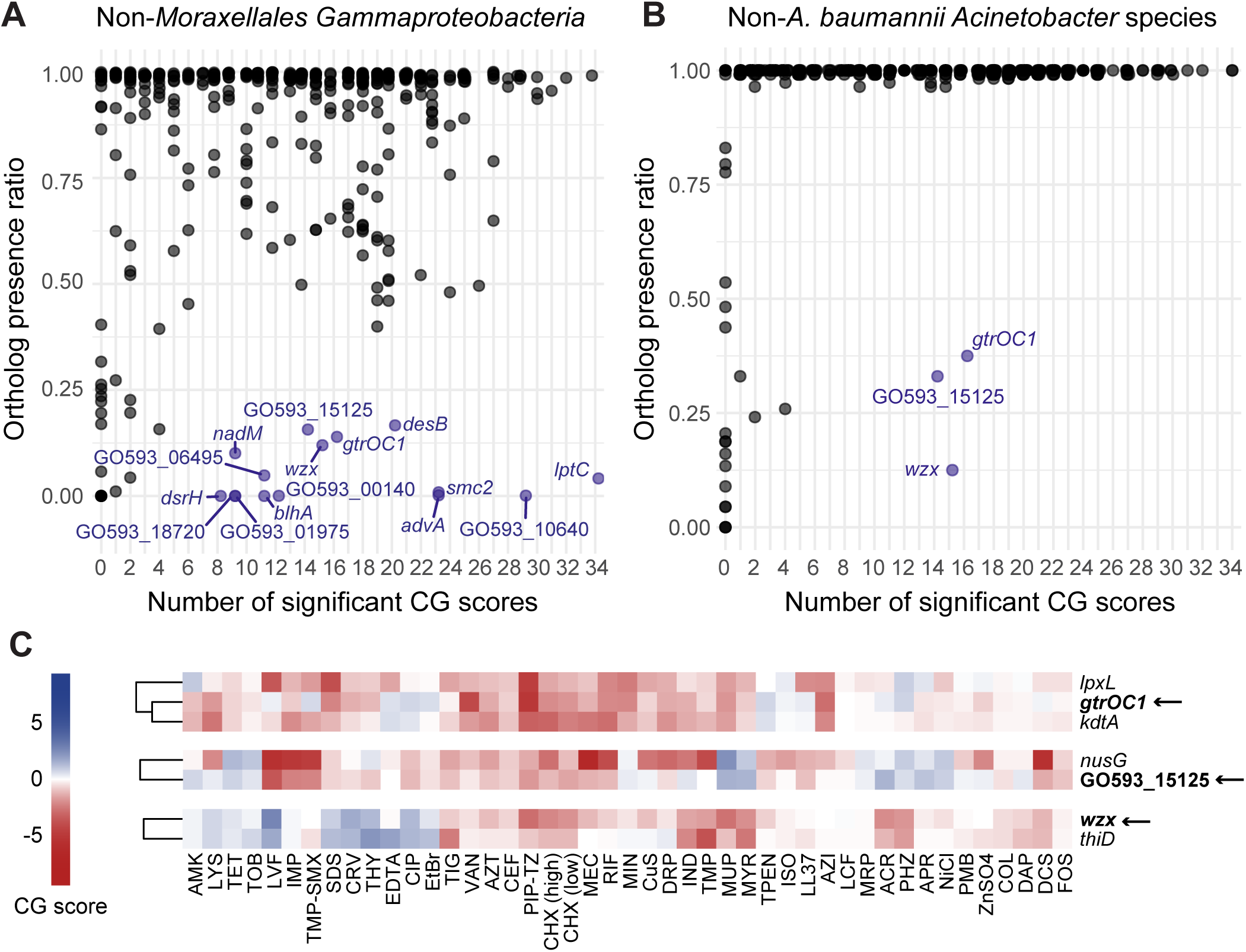
Chemical interactions and conservation of essential genes. (A) Dot plot depicting library chemical-gene scores and ortholog presence ratio (fraction of isolates possessing at least one ortholog out of the total number of analyzed isolates) across (A) representative non-*Moraxellales Gammaproteobacteria* or (B) non-*A. baumannii Acinetobacter* species. Dots in blue represent genes with <38% presence across representative groups and significant chemical-gene interactions in >10% of screen conditions. (C) Heatmap depicting CG scores for knockdowns of hits for genes rare outside *A. baumannii*.

Among others, this revealed that compared to other *lpt* genes orthologs to *lptC*, which is involved in lipooligosaccharide transport, are considerably rare among *Gammaproteobacteria* (Fig 3A; Appendix Fig S4). Additional underexplored cell envelope genes—GO593_05760 (*advA*), GO593_07665 (*blhA*), and GO593_11915 (*smc2*), GO593_03530 (*gtrOC1*), GO593_06865 (*wzx*),—suggest distinct and under-characterized components of the *A. baumannii* cell envelope may play key roles in chemical response.

We then increased our resolution to pinpoint genes specific to *A. baumannii* compared to other *Acinetobacter* species. Three genes stood out as more responsive to chemicals upon knockdown: *gtrOC1, wzx,* and GO593_15125 (Fig 3B). Both *gtrOC1* and *wzx* are involved in cell surface processes; *wzx* encodes a predicted polysaccharide flippase within a capsule biosynthesis locus (31), while *gtrOC1* is part of a locus involved in LOS synthesis (32). Orthologs to these genes are only sporadically found in *Gammaproteobacteria* outside *A. baumannii* (Fig 3A and B; Appendix Fig S3). For *wzx,* one explanation for this patchy occurrence of orthologs is the diversity of polysaccharides that the encoded protein transports (33, 34).

Similarly, *gtrOC1* has been implicated in a novel mechanism for core oligosaccharide assembly in *A. baumannii* (32). Hierarchical clustering of gene knockdowns based on CG score correlations using Ward’s method confirmed association of *gtrOC1* with LOS synthesis genes but revealed an unexpected clustering of *wzx* with *thiD*, a thiamin synthesis gene, for unknown reasons (Fig 3C).

GO593_15125 shows a similar ortholog distribution (Fig 3A and B; Appendix Fig S3) but encodes for an Arc-family repressor. In our previous work, we showed knockdown of a different Arc-family repressor caused significant fitness loss, likely due to its role in repressing toxic prophage genes (20). GO593_15125 could play a similar role in regulating mobile genetic elements (MGEs) as it is linked to a predicted transposase and an antiphage defense island (35, 36)). Additionally, the knockdown is most sensitive to levofloxacin, a DNA-damaging antibiotic (Fig 3C), and MGEs often modulate expression in response to DNA damage (37, 38) and occasionally trigger DNA damage responses themselves (39). Furthermore, this gene clusters with *nusG*, encoding a transcription elongation/termination factor (Fig 3C). NusG in *E. coli* induces the DNA damage response and *rac* prophage excision when depleted (40) and the essentiality of *nusG* is tied to suppression of toxic genes in *rac* (41), suggesting that GO593_15125 and *nusG* are both involved in prophage gene repression in *A. baumannii*. These findings identify key chemically responsive genes in *A. baumannii*, which are less common in other Gammaproteobacteria. Their potential unique roles in bacterial survival provide a strong foundation for further investigation into their contribution to drug resistance.

### An essential gene network in A. baumannii identifies novel gene connections

We next sought to identify further functional connections for these taxon-specific, chemically responsive genes. Using the CG scores, we calculated all pairwise correlations between genes then defined a significance threshold by generating an empirical null distribution through random permutation of the underlying data (7, 16). We used only correlations passing the significance threshold to construct a high-confidence essential gene network. Most connections in this network were between genes in different operons (Fig 4A), suggesting true functional relationships dominated over artifacts of CRISPRi polarity. As expected, genes in well-conserved pathways like translation and cell division showed numerous connections (Fig 4A). However, oxidative phosphorylation complexes were disconnected (Fig 4A; Appendix Fig S5A), suggesting distinct functionalities outside of the electron transport chain (e.g., succinate dehydrogenase also functions in the TCA cycle).

**Figure 4.**
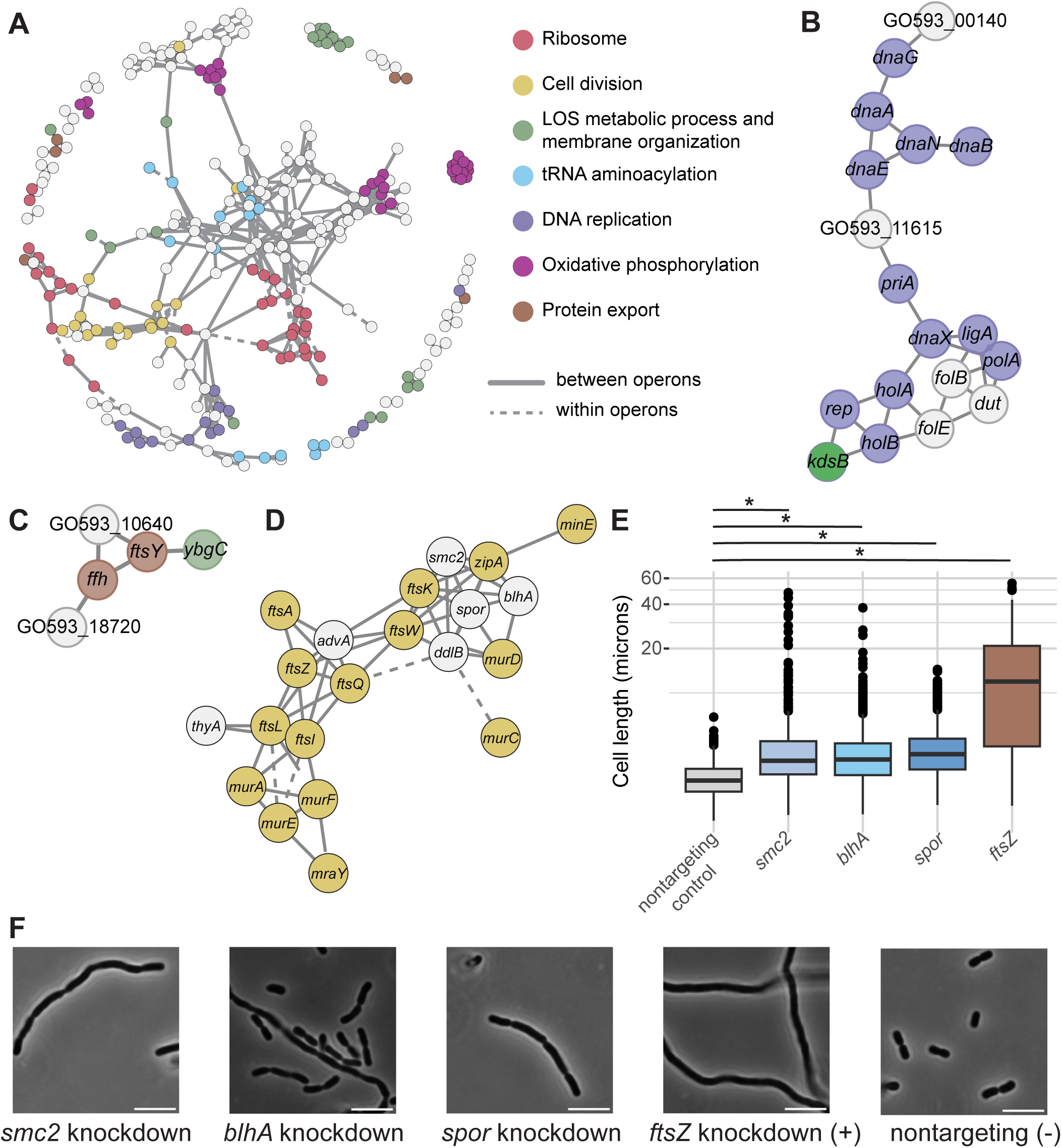
(A) Essential gene network in *A. baumannii*. Nodes (genes) are connected by edges (phenotypic correlations, r>0.76). Solid lines represent genes across different operons; dotted lines represent genes within the same operon. Subnetworks depict (B) DNA polymerase and replication-related genes, (C) signal-recognition, and (D) cell division and cell wall synthesis. (E) Quantification of cell length of knockdowns from microscopy of cell wall gene knockdowns (N=200-400). (F) Representative microscopy images. Ruler is 5 microns.

Connections in this network between poorly understood, taxon-specific genes and well-characterized, conserved genes point to shared functions. Lpt genes formed a subnetwork with *gtrOC1*, supporting the gene’s role in LOS synthesis (Appendix Fig S5B). In comparison, *wzx* was located in a subnetwork with genes involved in thiamin synthesis, ferredoxins, iron-sulfur cluster biogenesis, isoprenoid synthesis, cytochrome *bo_3_*oxidase, and another monovalent cation/H+ antiporter, hinting at broader connections between aerobic metabolism and isoprenoid lipid-linked glycans operated on by Wzx (42) (Appendix Fig S5A).

One of the candidate genes restricted to *Moraxellales*, GO593_00140, was linked with multiple DNA replication genes and an unannotated gene, GO593_11615 (Fig 4B). Knockdowns of GO593_00140 and GO593_11615 showed significant, negative CG scores in levofloxacin, which targets replication (Fig EV2A and B). Despite poor conservation (Fig 2A; Appendix Fig S3), structural prediction of GO593_00140 using Phyre2 showed a domain similar to that of *E. coli* HolD, the psi subunit of the bacterial DNA polymerase III holoenzyme (48% of the sequence modelled with 96.8% confidence despite only 12% sequence ID) (Fig EV2C). A previous hidden Markov model (HMM)-based homolog search identified similar putative psi subunits in other *Acinetobacter* species (43), and our work provides phenotypic evidence to further support that this gene is involved in DNA replication in *A. baumannii*. GO593_11615 likely encodes a phage-associated transcriptional regulator, as the gene is situated within an intact prophage predicted by PHASTEST (44). This protein has domain architecture similar to LexA, a conserved repressor of DNA repair pathways that senses DNA damage through an interaction with RecA-DNA filaments that triggers self-cleavage and subsequent de-repression of genes involved in the SOS response (45) (Fig EV2D). Depletion of LexA is known to induce excision of some prophages and subsequent host lysis (37). Interestingly, *A. baumannii* lacks *lexA* (46), suggesting this prophage encodes its own LexA-like protein as a host-independent strategy to detect DNA damage.

Other candidate genes, GO593_10640 and GO593_18720, grouped with the conserved signal recognition particle (SRP) genes *ftsY* and *ffh* as well as *ybgC*, a Tol-Pal-associated thioesterase linked to membrane integrity (47) (Fig 4C). GO593_10640 shows structural similarities to ion transporters and significant chemical responses consistent with *ftsY* and *ffh* knockdowns (Appendix Fig S6A and B). GO593_18720, however, showed CG scores that were smaller in magnitude, suggesting a less critical or defined role (Appendix Fig S6A). These genes may be directly or indirectly related to membrane protein localization and membrane integrity, though SRP mechanisms remain understudied in *A. baumannii*.

Our network analysis revealed a cell wall and division subnetwork (Fig 4D), containing three genes we identified as rare in other *Gammaproteobacteria*: GO593_05760 (*advA*), GO593_07665 (*blhA*), and GO593_11915 (*smc2*, or *SMC_prok_B*). These genes correlate with numerous well-characterized, highly conserved division genes, corroborating initial data from other studies suggesting roles for *advA*, *blhA*, and *smc2* in cell division (3, 48). This subnetwork also featured another gene of unknown function, GO593_07290, which encodes for a SPOR domain-containing protein, annotated here as *spor*. While SPOR domains are widely conserved and are known to interact with cell wall peptidoglycan (PG) (49), functions of SPOR domain-containing proteins vary considerably. Indeed, one of the most extensively characterized SPOR domain proteins, FtsN from *E. coli*, does not require the SPOR domain for its essential function (50). The short periplasmic region required for FtsN function, ^E^FtsN (51), does not appear to be present at the amino acid level in GO593_07290, suggesting a functionally distinct role in cell division (Fig EV2E).

To further support these genes’ roles in cell division, we generated individual CRISPRi knockdowns for *blhA*, *smc2*, and *spor* and examined cell shape with microscopy. Knockdown of *advA* has previously been shown to cause filamentation, a hallmark of impaired division (12). Similarly, *blhA, smc2*, and *spor* knockdowns produced elongated or chained cells (Fig 4E-F), corroborating prior findings from a *blhA* transposon mutant (48). We next attempted to determine the localization pattern of putative division genes using *trans* expression of fluorescent protein fusions. Similar to what had been previously shown with AdvA (3), a fusion of superfolder GFP to the N-terminus of BlhA (sfGFP::BlhA) showed localization to the septum, or the site of division, suggesting that BlhA is a bona-fide member of the divisome (Fig EV3A). In contrast, the SPOR domain protein localized to both the cell periphery and septum (Fig EV3B), consistent with PG binding, but suggests a more generalized cell wall synthesis function.

Interestingly, expression of the Smc2::sfGFP hybrid resulted in cell elongation and division defects, with the fluorescently tagged Smc2 protein forming elongated aggregates in the cells (Fig EV3C). The impaired division phenotypes were verified with overexpression of Smc2 alone, suggesting precise protein levels are needed for proper cell division (Fig EV3D). This *trans* expression additionally allowed us to disentangle the role of *smc2* from polar knockdown effects on downstream gene *zapA*, a known division gene. As an important caveat, we note that our fusion constructs may not be expressed at the same level as their native counterparts and, therefore, may show aberrant localization. Nonetheless, we contend that these localizations are at least possible. Overall, our network reveals functional connections and underlying phenotypes linking crucial pathways in *A. baumannii*, such as cell division, and provides a basis for further targeted experiments for mechanistic analysis.

### Structure-informed clustering of chemical perturbations reveals insights into inhibitor function

We next aimed to develop an approach for characterizing inhibitor function in *A. baumannii* that leveraged our chemical genomics dataset. To broadly define inhibitor impacts on cell physiology, we analyzed chemical-gene interactions at the pathway level rather than the individual gene level. This approach evaluated the L_2_FCs of all guides, including mismatches, associated with a STRING functional pathway (“pathway effect”) (52), enhancing signal robustness. We then clustered chemicals by pathway-level interactions, showing the relationship between chemicals solely based on the cellular pathways they impacted in our screen (Fig 5A). Inhibitors from the same antibiotic classes, such as quinolones (ciprofloxacin and levofloxacin) or aminoglycosides (amikacin, apramycin, and tobramycin), clustered as expected, validating our method.

**Figure 5.**
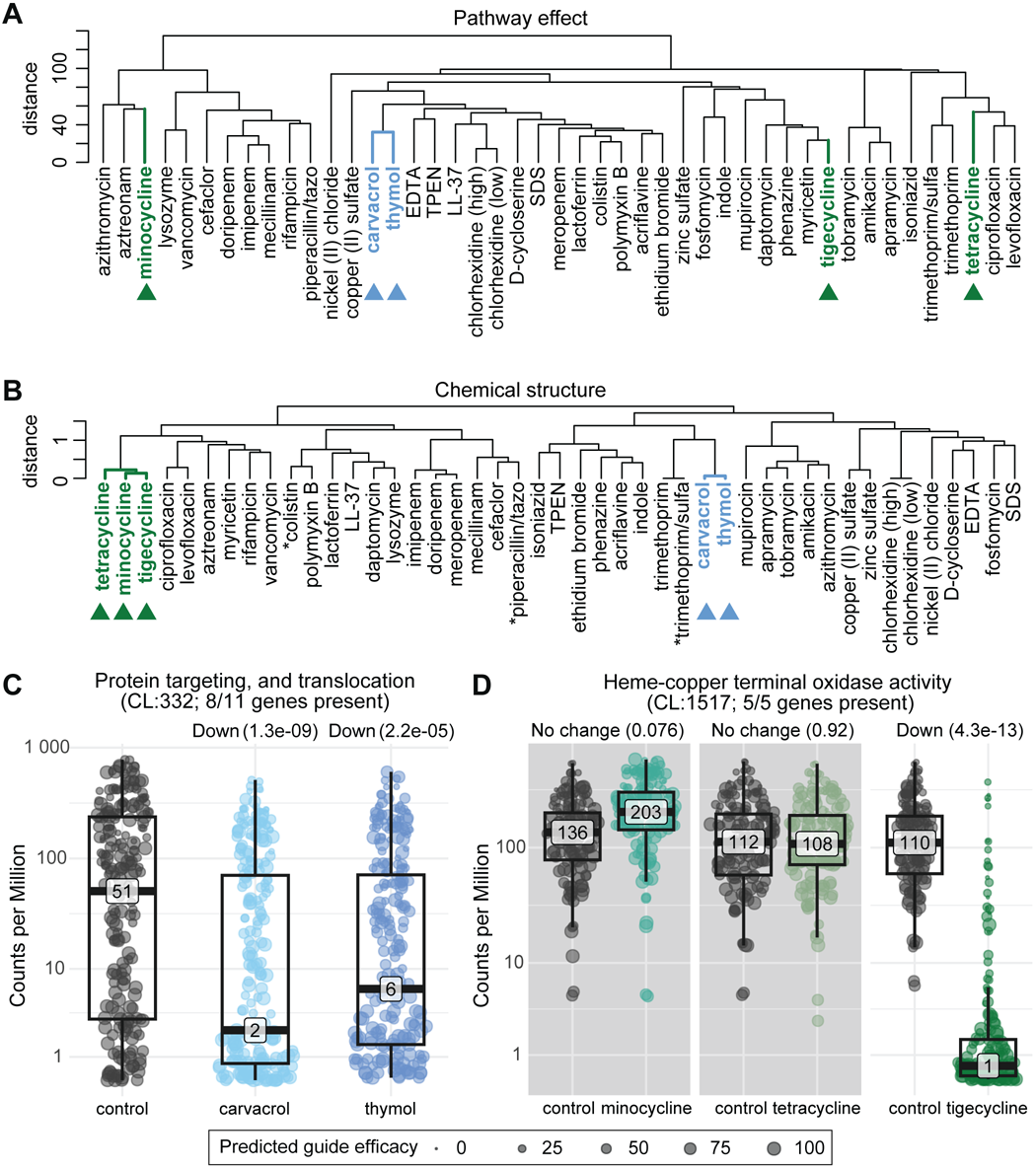
Chemical grouping by pathway effects and structure. (A) Dendrogram depicting hierarchical clustering (Ward method) of screen chemicals by similarities of effects across essential gene pathways. (B) Dendrogram displaying hierarchical clustering (Ward method) of chemical molecular fingerprints (ECFP6) by Tanimoto coefficient. Asterisks represent conditions with multiple chemical structures, in which a single structure was used (see Table S4). (C) Sina plots depicting differential guide counts per million (CPMs) for protein translocation in carvacrol and thymol, or (D) for heme-copper terminal oxidase in tetracycline-class antibiotics, compared to corresponding no chemical controls. Chemical-gene interactions are described above each graph—up (positive), down (negative), or no change—with FDR values. Guides are weighted in this calculation by predicted efficacy, with perfect guides weighted at 100. Non-significant comparisons are shown with grey backgrounds.

To more systematically compare our results to inhibitors outside of established antibiotic classes, we performed molecular fingerprinting—a chemoinformatic approach that numerically represents unique chemical structure and properties of molecules as binary “fingerprints” (extended-connectivity fingerprints) (53, 54)—and measured similarity between chemicals using the Tanimoto coefficient (55, 56). Clustering based on distances derived from Tanimoto coefficients verified that antibiotics with similar core structures (beta-lactams, fluoroquinolones, aminoglycosides, etc.) group together (Fig 5B). Comparing the pathway impact clustering with the structure clustering of these chemicals allowed us to investigate (a) consistent chemical classes with a not-yet-established mechanism of action, (b) structurally similar chemicals that have distinct effects on *A. baumannii*, and (c) structurally dissimilar chemicals that similarly affect *A. baumannii* physiology.

One example is carvacrol and thymol, which cluster tightly by both pathway effect and structure (Fig 5A and B; Appendix Fig S7). These compounds exhibit antibacterial activity but lack a defined mechanism of action (57), though they have been proposed as antimicrobial agents or additives (58). Our pathway analysis revealed NADH dehydrogenase (*nuo* genes) and protein transport (*ftsY, ffh,* and *sec* genes) knockdowns were significantly sensitized while ubiquinone (*ubi* genes) knockdowns were more resistant (Fig 5C and EV4A-D). In particular, *ftsY*, encoding the SRP receptor protein, was a major negative outlier in both carvacrol and thymol (Fig EV4C-D). These pathways influence membrane homeostasis and proton motive force (PMF), supporting previous studies stating that membrane disruption and depolarization underlie carvacrol and thymol antibacterial activity (59). However, carvacrol and thymol cluster more distantly in their pathway effects from other known membrane disruptors, like chlorhexidine and EDTA (60, 61) (Fig 5A). This suggests that the mechanisms of action for these phenolic monoterpenes are unique from may be unique, potentially due to more specific effects on SRP-mediated protein transport.

Identification of structurally similar chemicals with differing effects on *A. baumannii* pathways could reveal how relatively small chemical modifications alter their physiological impact. To explore this, we scored how well chemical pairs clustered in pathway effect or in structure and calculated the differences (Appendix Fig S8). Notably, the pathway effects of tetracycline, minocycline, and tigecycline diverged despite their strong structural similarity (Fig EV5C).

Surprisingly, knockdowns of ribosomal genes as a whole did not show significant differences in abundance when treated with tetracycline-class antibiotics (Appendix Fig S9A-B). Ribosomal genes lacked significant CG scores across ribosome genes in minocycline or tetracycline— potentially due to efflux mechanisms or other resistance factors impacting efficacy—and showed high variability in significant CG scores across ribosome genes for tigecycline, suggesting variability between ribosomal components (Appendix Fig S9C). Knockdowns in the LOS transport pathway, Lpt, generally increased sensitivity to these antibiotics, although the extent and confidence of the pathway effect varied (Fig EV5A). The most significant divergences in pathway effect of these drugs were pathways contributing to membrane potential and oxidative phosphorylation. ATP synthase knockdowns exhibited increased abundance in tigecycline and tetracycline, but not minocycline (Fig EV5B). RND efflux pumps in *A. baumannii* are often able to efflux tigecycline and tetracycline but not minocycline (62). However, the relationship between decreased RND-mediated efflux and ATP synthase knockdowns remains unclear, especially as compensatory mechanisms could confound observed effects. Unlike other tetracyclines, tigecycline contains a unique structural modification making it a glycylcycline (63). This structural change may explain tigecycline’s CG score variation for ribosomal genes and its unique effectiveness against heme-copper terminal oxidase (cytochrome *bo_3_*oxidase) knockdowns (Fig 5D) despite the shared core structure with minocycline and tetracycline. Based on pathway effect, tigecycline clusters with the structurally dissimilar chemicals phenazine and myricetin. While the exact mechanisms of action of phenazine and myricetin are unclear, both are known to affect oxidative phosphorylation and oxidative stress (64–67), suggesting tigecycline may similarly affect oxidative phosphorylation in *A. baumannii*.

We note that chemicals may display reduced pathway impacts in this analysis due to ineffective or noncomparable concentrations used and result in apparent divergence from their structural groups, as was likely the case with meropenem (vs other carbapenems) in this screen (Appendix Fig S10). Overall, we established a systematic method to group and compare chemicals by genetic interactions and structure, successfully highlighting effects of an understudied chemical group and uncovering unique effects of tetracycline-class antibiotics on *A. baumannii*.

## Discussion

Essential cellular processes are the current and likely future targets of mono- and poly-therapies against antibiotic resistant pathogens. Despite the biological and clinical importance of essential genes, few studies have comprehensively characterized their functions, especially in Gram-negative pathogens. This work advances our understanding of gene and antibiotic function in *A. baumannii* through large-scale, chemical-genomic profiling of an essential gene knockdown library. The breadth and diversity of our chemical perturbations provided functional insights that could not be obtained with a smaller scope. We identified >5000 significant phenotypes for essential genes (Fig. 1) and found that specific pathways, such as LOS transport, were highly enriched for sensitive phenotypes (Fig. 2), representing compelling targets for combination therapies. Our evolutionary analysis revealed inhibitor-responsive essential genes that are unique to or are highly divergent in *A. baumannii* (Fig. 3), providing possible targets for narrow-spectrum therapeutics. Our essential gene network showed robust connections between known and poorly characterized essential genes (Fig. 4), hinting at new functions and corroborating prior work. Finally, our structure-function analysis of chemical-gene phenotypes uncovered a surprising discordance among the phenotypic impacts of antibiotics within the same class (Fig. 5), highlighting how genes outside of the direct target can alter antibiotic efficacy. Moreover, this study provides a rich dataset of essential gene phenotypes to the *A. baumannii* and larger Gram-negative pathogen research communities, serving as a resource and framework for furthering our understanding of essential gene function.

Our findings linking Lpt function to antibiotic resistance accentuate the potential of targeting LOS trafficking for combination therapies, especially in the context of recent pharmaceutical advances.

Our observation that *lpt* knockdown strains show a hyper-permeable phenotype that is dependent on LOS production suggests that increased permeability is caused by both loss of LOS in the outer membrane and retention of LOS in the inner membrane, although additional work is needed to show this definitively. Other studies have suggested that inhibiting LPS transport in other Gram-negatives results in toxic accumulation of LOS intermediates that cause membrane defects and antibiotic susceptibility (68–70); these hypotheses are not mutually exclusive with our own. Moreover, the increased permeability of *lpt* knockdowns is consistent with classic *E. coli lptD* (*imp4213* (71)) mutants that are broadly permeable (72) and are often used in drug screening efforts (73). Recent studies have characterized novel Lpt-targeting macrocyclic peptides (74, 75) that selectively disrupt LOS transport in *Acinetobacter* species.

Interestingly, our conservation analysis revealed that LptC is far more divergent among Gammaproteobacteria than other proteins in the Lpt complex, which may be related to the fact that these Lpt-targeting macrocyclic peptides are inactive against other Gram-negatives. Taken together, these data highlight the potential for recently described LOS transport inhibitors (70, 74, 75) to be effective in conjunction with other antibiotics as a combination therapy against *A. baumannii* infections.

Our work also raises intriguing questions regarding the functions of essential genes common in *A. baumannii* yet rare across other Gammaproteobacteria. Our essential gene network revealed functional connections between many of these poorly characterized and well-studied ones, generating hypotheses that can be validated in future mechanistic studies. For instance, GO593_10640 knockdown strains show strong phenotypic similarity to knockdowns of SRP components. Whether GO593_10640 is required for SRP activity or is a key SRP substrate involved in antibiotic resistance is unknown. GO593_10640 contains a predicted signal sequence, and, thus, is a potential SRP substrate. SRP is poorly characterized in *A. baumannii*, although recent work has highlighted its potential as an antimicrobial target (76). Advances in protein structural modeling have caused a paradigm shift in contemporary biology (77); synthesizing structural predictions with systematic phenotyping is becoming a core strategy to elucidate gene functions. Using this approach, we uncovered GO593_00140 as a possible *holD* analog in *A. baumannii*. Although GO593_00140 is highly divergent from *E. coli* HolD at the primary sequence level, it contains similar predicted folds and a GO593_00140 knockdown strain has overlapping chemical phenotypes to knockdowns of other DNA polymerase III (DNAP) subunits, corroborating previous in silico predictions (43). Additionally, we identified a possible LexA analog (GO593_11615) inside a predicted prophage. Although *A. baumannii* has a distinct DNA damage response that does not rely on LexA (46), knockdown strains of this analog display phenotypes that match disruption of DNAP activity. These findings open the possibility that mobile genetic elements could serve as a resource to better understand the DNA damage response or identify proteins that are particularly toxic to *A. baumannii*. Finally, our ortholog and network analyses suggest that *A. baumannii* contains may unique or divergent genes associated with cell division. Synthesizing our work with prior studies (3, 48) supports the hypothesis that *A. baumannii* contains a unique divisome with AdvA and BlhA localizing to the division septum and additional proteins possibly participating in division either directly (SPOR) or indirectly (Smc2). The precise functions of these players and their order of assembly into the divisome (78) has yet to be elucidated.

Our structure-function studies combining chemoinformatics, bioinformatics, and phenotype data provided insights into the functions of known and poorly characterized inhibitors. Our results with carvacrol and thymol, two inhibitors of unknown mechanism, suggest that perturbance of membrane associated complexes such as NADH dehydrogenase complex I and the SRP underly their toxicity to *A. baumannii*. This is broadly consistent with previous studies that have noted that thymol disrupts PMF (59) or generates reactive oxygen species through the Fenton reaction (79). The direct target remains unknown; indeed, multiple targets are possible. We further show that antibiotics within the same class can have divergent effects on *A. baumannii* physiology. Among tetracycline class antibiotics, only minocycline was unaffected by ATP synthase knockdowns and only tigecycline showed increased efficacy against cytochrome *bo_3_* knockdowns. Despite tigecycline being a more recently synthesized tetracycline derivative, it is often less effective against *A. baumannii* than minocycline due to efflux (62). Efflux is dependent on the cellular energy state (80), which may link the cytochrome *bo_3_* oxidase to reduced tigecycline efficacy. Previous work suggests that tigecycline is more prone to inactivation by dissolved oxygen in media than other tetracyclines (81) and is known to have anti-cancer activities due to induction of mitochondrial oxidative stress (82). Cytochrome *bo_3_* oxidase, as the main terminal oxidase in *A. baumannii*, has an important role in reducing oxygen that could impact tigecycline stability. It is important to note that these genes are essential in *A. baumannii* because of its obligate aerobic lifestyle, and knockdowns of ATP synthase or cytochrome *bo_3_* oxidase may be causing complex compensation or stress responses that affect observed phenotypes. Our identification of distinct physiological impacts of tetracyclines underscores the utility of our chemical-genomics approach to identify mechanistic differences between closely related compounds.

Overall, this study enhances our understanding of essential genes in *A. baumannii*, highlighting unique aspects of its biology and establishing a useful resource for future studies. A limitation of our approach is dependence on CRISPRi, which causes polar effects on downstream genes in operons and can have off-target effects, although our experimental design and analysis strategy mitigate some of the risks of toxic or otherwise off-target guides. Additionally, much of our computational analysis relies on similarity to known proteins and functional predictions, particularly from the STRING database (24). *A. baumannii* has a poorly characterized proteome compared to model bacteria; despite this, our approach still successfully identified many expected gene connections, likely due to the high level of conservation in essential gene sequences and functions. Future studies could integrate our dataset or network analyses to enhance the breadth and coverage of predicted genetic pathways. While this screen focuses on essential genes and a curate set of chemicals known to cause growth defects in *A. baumannii*, our systems approach can be readily applied to full knockdown libraries or small molecule screens in other pathogens.

## Methods

### Bacterial strains and growth

All strains, plasmids, and oligos used in this study are listed in Tables S1-3. Strains were grown in EZ Rich Defined Media (Teknova) supplemented 40 mM sodium succinate as the carbon source (AbRDM) or AbRDM + 1.5% agar at 37°C unless otherwise noted. Selective antibiotics were used when necessary: for *E. coli*, 100 µg/mL ampicillin (amp) or carbenicillin (carb), 15 µg/mL gentamicin (gent), or 30 µg/mL kanamycin (kan); and for *A. baumannii*, 10 µg/mL polymyxin B (pmb), 150 µg/mL gentamicin (gent), 60 µg/mL kanamycin (kan). Diaminopimelic acid (DAP) was added at 300 µM to support growth of E. coli dap-donor strains. IPTG (isopropyl b-D-1-thiogalactopyranoside) (0 to 1 mM) was added where indicated in the figures or figure legends. Strains were preserved in 15% glycerol at −80°C.

### General molecular biology techniques and plasmid construction

Plasmids, oligos, and other construction details are listed in Tables S2-3. Oligonucleotides were synthesized by Integrated DNA Technologies (Coralville, IA). Plasmid DNA was purified using the GeneJet plasmid miniprep kit (K0503; Thermo Scientific); genomic DNA was purified using the GeneJet genomic DNA purification kit (K0721; Thermo Scientific). DNA fragments were amplified with Q5 DNA polymerase (M0491; New England Biolabs (NEB)) or OneTaq DNA Polymerase (NEB). All enzymes for restriction digests were from NEB. DNA fragments were spin-purified using DNA Clean & Concentrator kit (D4004; Zymo Research) after digestion or amplification. Ligations performed using T4 DNA ligase (M0202; NEB). Plasmids were assembled from restriction enzyme-linearized or PCR-amplified vector and PCR products or synthetic DNA fragments using the NEBuilder Hifi DNA assembly kit (E2621; NEB). Plasmids were transformed into electrocompetent E. coli cells using a Bio-Rad Gene Pulser Xcell on the EC1 setting. Sanger DNA sequencing was performed by Functional Biosciences (Madison, WI); full plasmid sequencing was performed by Plasmidsaurus.

### Library growth experiment

The *A. baumannii* essential gene CRISPRi library (sJMP2949) was diluted 1000-fold from frozen stock (OD600=15) in 50 mL AbRDM (starting OD600= 0.015) and incubated in 250 mL flasks at 37°C with shaking until mid-log (OD600=0.2; ∼3.5 hrs)(t0). This culture was then diluted to OD600 =0.02 in 4 mL media with 1mM IPTG and antibiotics (antibiotic stocks diluted 1:100 into media and used immediately; see Table S4) in 14 mL snapcap culture tubes (Falcon 352051) and incubated with shaking at 37°C. At 18 hours, saturated cultures were serially diluted back to OD600=0.02 into fresh tubes containing the same media and incubated for another 18 hours at 37°C with shaking, for a total of ∼14 doublings. Cells were pelleted from 1 mL culture in duplicate before (t0) and after growth in antibiotics.

### Sample preparation and sequencing

Genomic DNA was extracted from cell pellets and resuspended in a final volume of 200 μL. The sgRNA-encoding region was amplified using Q5 DNA polymerase (NEB) in a 100 μL reaction with ∼100 ng gDNA and primers oJMP697 and oJMP698 according to manufacturer’s protocol, using a BioRad C1000 thermocycler: 30s at 98°C, followed by 16 cycles of 15s at 98°C, 15s at 65°C, 15s at 72°C, then 72°C for 5 min. PCR products were spin-purified and eluted in a final volume of 20 μL.

Samples were sequenced by the UW-Madison Biotech Center Next Gen DNA Sequencing Core facility. PCR products were amplified with nested primers containing i5 and i7 indexes and Illumina TruSeq adapters, cleaned, quantified, pooled, and run on a Novaseq 6000 (150 bp paired-end reads).

### Sequencing read quality control and processing

Guides were counted using a custom Python script designed to minimize noise and accurately count guides with overlapping targets. This script samples sequencing reads to identify barcode diversity, orientations, and offsets, and determines flanking sequences for correct barcode identification. Only counts for library barcodes were used in downstream analyses.

Coefficients of determination (R²) between samples were calculated from barcode counts per million (CPM), and conditions with replicates where R² < 0.5 were removed to ensure replicate agreement. Samples were also excluded if population diversity (Nb) was less than 10,000 for nontargeting control guides.

Log_2_ fold changes and confidence values were computed using edgeR. Gene-level values were calculated by taking the median guide-level log_2_ fold change for perfect match guides; confidence was determined by computing Stouffer’s p-value using guide-level FDR values (poolr R package). Functional enrichment analyses were conducted using STRING-db version 11.5 with a custom *A. baumannii* ATCC 19606 proteome (Organism ID: STRG0060QIE).

### Ortholog analyses

Genome assemblies of *Gammaproteobacteria* were collected from the NCBI Reference Sequence database (RefSeq release 213). The dataset comprises 2792 isolates distributed among 474 genera and 18 taxonomic orders (Table S8).

Essential genes represented in the library were pulled from the genome assembly for *A. baumannii* ATCC 19606 (GCA_009759685.1). Orthology assignment was computed with the phylogenetic profiling tool fDOG v.0.1.26 (30), compiling a maximum of 25 core-orthologs for each gene using a taxonomic distance minimum of genus and maximum of class. Visualizations were performed using PhyloProfile (83).

The ortholog presence ratio for each gene was measured within a taxonomic group by calculating the fraction of isolates possessing at least one ortholog out of the total number of analyzed isolates. We evaluated these ratios at the genus level with non-ACB (*Acinetobacter calcoaceticus-baumannii* complex) isolates, at the order level with non-*Acinetobacter Moraxellales*, and at the class level with non-*Moraxellales Gammaproteobacteria*.

### Creation of knockdown strains

As previously described (20, 84), sgRNA-encoding sequences were cloned between the BsaI restriction sites of MCi plasmid pJMP2776. Two oligonucleotides encoding an sgRNA were designed to overlap such that when annealed, their ends were complementary to the BsaI-cut ends on the vector.

Next, the Mobile-CRISPRi system was transferred to the attTn*7* site on the chromosome of *A. baumannii* by quad-parental conjugation of three donor strains - one with a mobilizable plasmid (pTn*7*C1) encoding Tn*7* transposase, another with a conjugal helper plasmid (pEVS104), and a third with a mobilizable plasmid containing a Tn*7* transposon encoding the CRISPRi system - and the recipient strain *A. baumannii* ATCC 19606 or ATCC 17978. Briefly, 100 µL of culture of donor and recipient strains were added to 600 µL LB, pelleted at ∼8,000 × g, washed twice with LB prior to depositing cells on a nitrocellulose filter (Millipore HAWP02500) on an LB plate, and incubated at 37°C, ∼4 h. Cells were removed from the filter by vortexing in 500 μL PBS and selected on LB-gent plates at 37°C.

### MIC test strip assays

500 μL of culture (OD=0.1, or 0.5 for LOS-strains) was spread on Mueller-Hinton agar (MHA) plates and allowed to dry for 30 minutes. A sterile MIC test strip was added to the plate and incubated overnight at 37°C. The MIC was determined as the value where the zone of inhibition meets the strip.

### Spot assays

Strains were grown on LB agar overnight. Cell scrapes were resuspended from plates into 500 μL PBS, and normalized to an OD_600_ of ∼1.0. 10 μL of ten-fold dilutions of these resuspensions were then spotted on AbRDM and AbRDM +1mM IPTG agar plates.

### Membrane permeability assay

ATCC 19606 and LOS-strains containing sgRNAs targeting *lptA* or a non-targeting control (sJMP4539 and 4324, respectively) were grown on LB agar or LB agar containing 1mM IPTG overnight. Cell scrapes were resuspended from plates into 500 μL PBS and normalized to an OD600 of 0.3. Ethidium bromide (EtBr; Bio-Rad #1610433) was added to cell suspensions at a final concentration of 10 ug/mL, and OD600 and fluorescence at 545nm excitation/600 emission were read in a plate reader (Tecan Infinite 200 Pro M Plex). Readings occurred over 100 cycles (∼75 min).

### Chemical comparison analyses

After edgeR was used to import, organize, filter, and normalize counts, the limma package with voom method was used to perform gene set testing using functional groups predicted by STRING-db (custom proteome; Organism ID: STRG0060QIE). Chemical effects on pathways were scored for direction and significance, then transformed into a correlation matrix. Chemicals were clustered using hierarchical clustering with the Ward method, and a dendrogram visualized the clustering based on pathway impacts.

SMILES strings for each chemical were parsed using the rcdk package in R to generate molecular fingerprints. For combination or mixed-structure antibiotics, a representative structure was selected (e.g., trimethoprim for trimethoprim-sulfamethoxazole or polymyxin E1 for colistin). Tanimoto coefficients were calculated to measure chemical similarity and generate a similarity matrix, which was then clustered using the Ward method to produce a chemical structure dendrogram.

To assess the relationships between chemical pairs, each dendrogram was iteratively cut at all k-values (1–45), and pairs were scored if they clustered together. The final association scores are the sums across all k-values. Differences in association scores between pathway-effect and structure were plotted to highlight disparities. Additionally, the sums of association scores for pathway-effect and structure were plotted to highlight similarities.

### Protein structural prediction

Structures of GO593_10640 and GO593_00140 (with chi subunit ortholog) were predicted from amino acid sequence using Alphafold2 (77, 85) and AlphaFold-Multimer (86). Predicted structures were visualized using PyMOL (Schrödinger).

### Cell division protein localization and microscopy

Expression vector plasmids were transferred by conjugation into *A. baumannii* ATCC 17978; briefly, 100 µL of culture of donor and recipient strains were mixed and deposited on a nitrocellulose filter (Millipore HAWP02500) on an LB plate, and incubated at 37°C, ∼2 h. Cells were removed from the filter by vortexing in 500 μL PBS and selected on LB-kan plates at 37°C.

Cultures were grown from a starting OD600∼0.01 to mid-log at 37°C for ∼3 hours in AbRDM + 1mM IPTG (or 500 uM IPTG for Smc2 and Smc2-sfGFP overexpression strains, due to toxicity). 1 mL of culture was centrifuged (8000xg for 2 min) and resuspended in equal volume PBS. Cells were fixed with paraformaldehyde (final concentration 5%) and quenched with equal volume 2.5M glycine. 10 μL of each sample were spotted on a glass slide for microscopy.

Bacteria were imaged with a Nikon Ti-E inverted microscope with an Orca Fusion BT digital CMOS camera (Hamamatsu) using NIS-Elements. Fluorescence images were collected using Prior Lumen 200 metal halide light source and a FITC-specific filter set. Cell length analysis was done with MicrobeJ (87).

## Data availability

The datasets and computer code produced in this study are available in the following databases:

- Raw sequencing data: NCBI BioProject (PRJNA1190454)
- Scripts: Github (https://github.com/jasonpeterslab)

## Supporting information

Extended view figures

## Acknowledgments

We thank the UW-Madison Biotech Center for their assistance with sequencing. Thanks to Stephen Trent and Brent Simpson for their gift of the LOS-strain, and special thanks to Rachel Salemi and Alex Goetsch for their assistance and training with microscopy. We thank Warren Rose and Tim Bugni for helpful comments. This work is supported by the National Institutes of Health under award numbers K22AI137122 and 1R35GM150487-01. JST is supported by the Biotechnology Training Program (NIH 5T32GM135066), SciMED Graduate Research Scholars, and an NSF GRFP. RDW is supported by the Predoctoral Training Program in Genetics (NIH 5T32GM007133) and SciMED Graduate Research Scholars.

