## Supplementary figures and images for "Chemical genomics informs antibiotic and essential gene function in *Acinetobacter baumannii*"

### Appendix figures

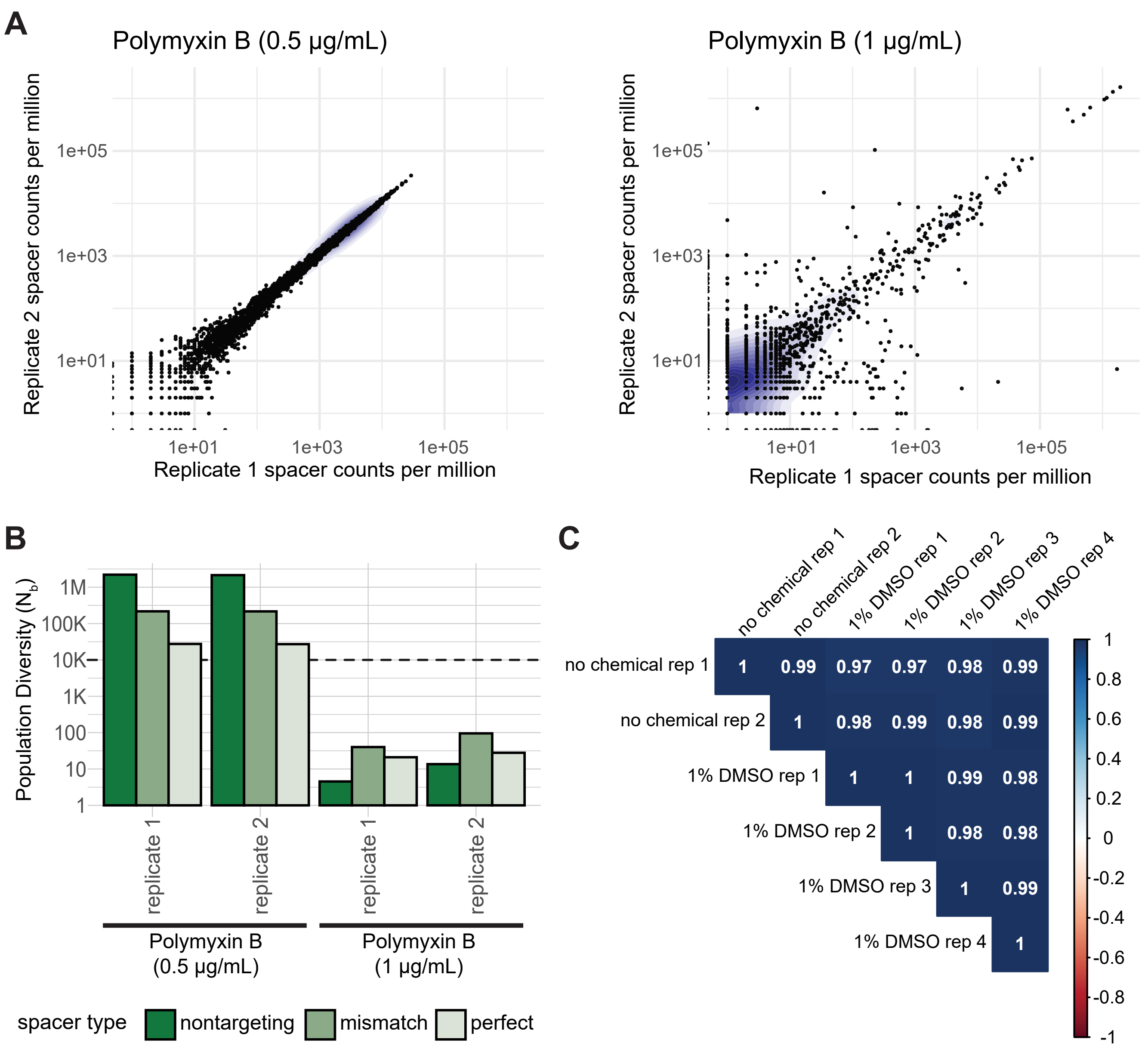

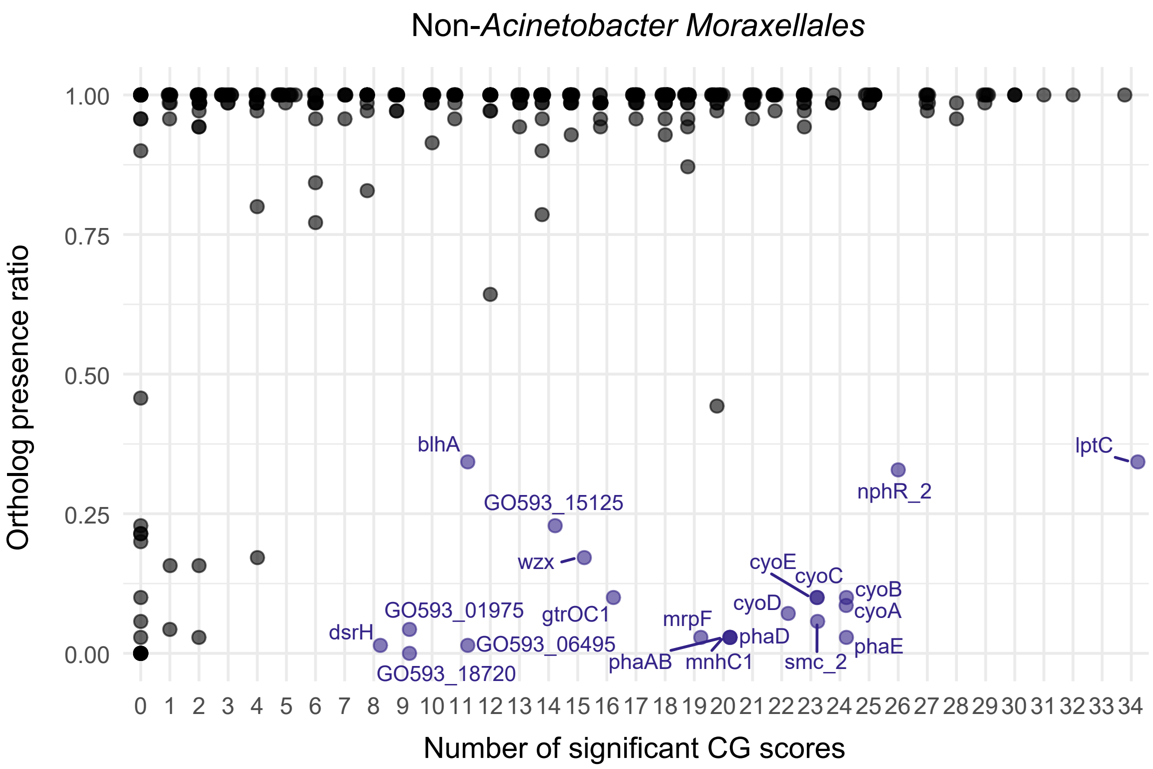

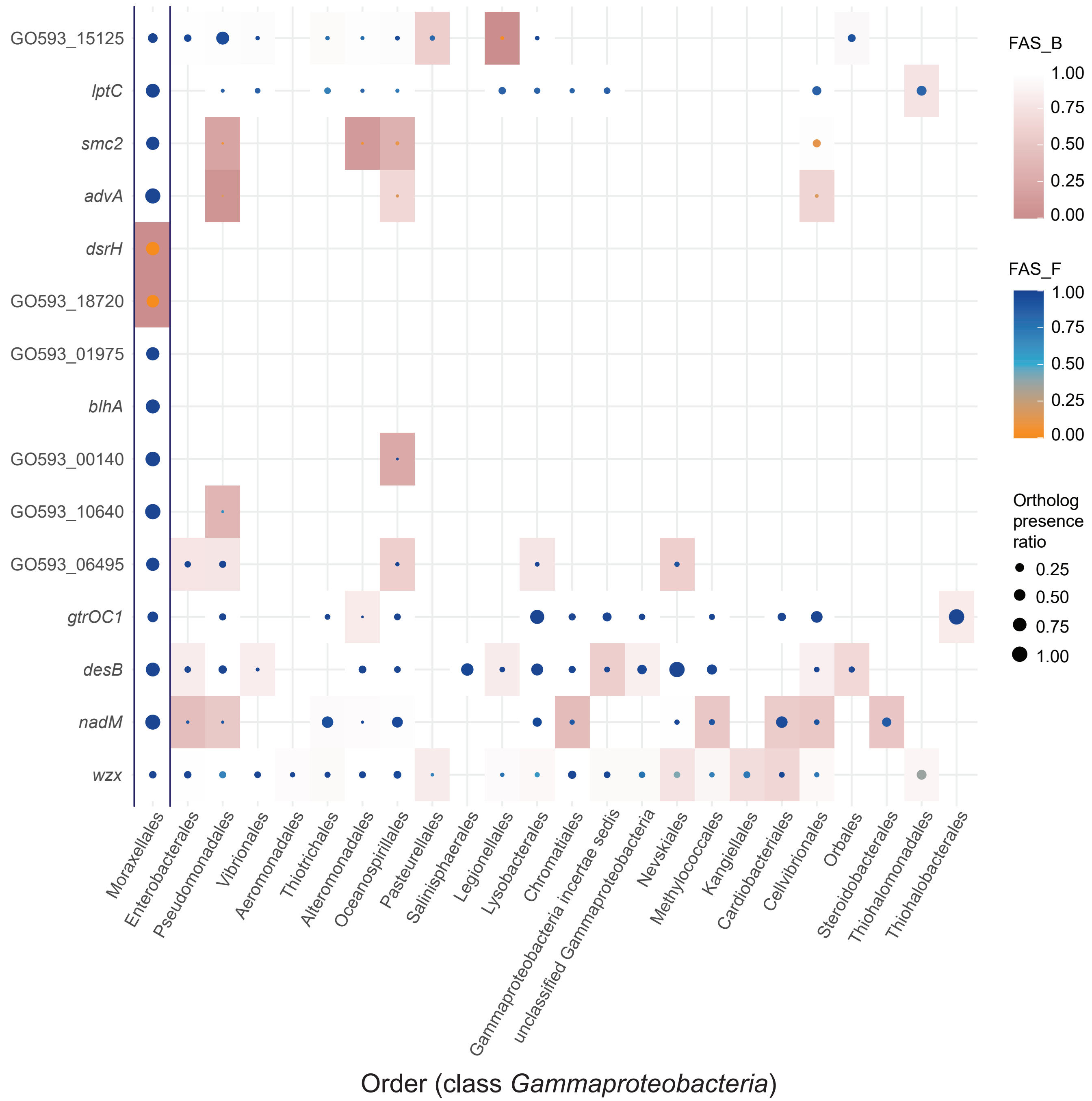

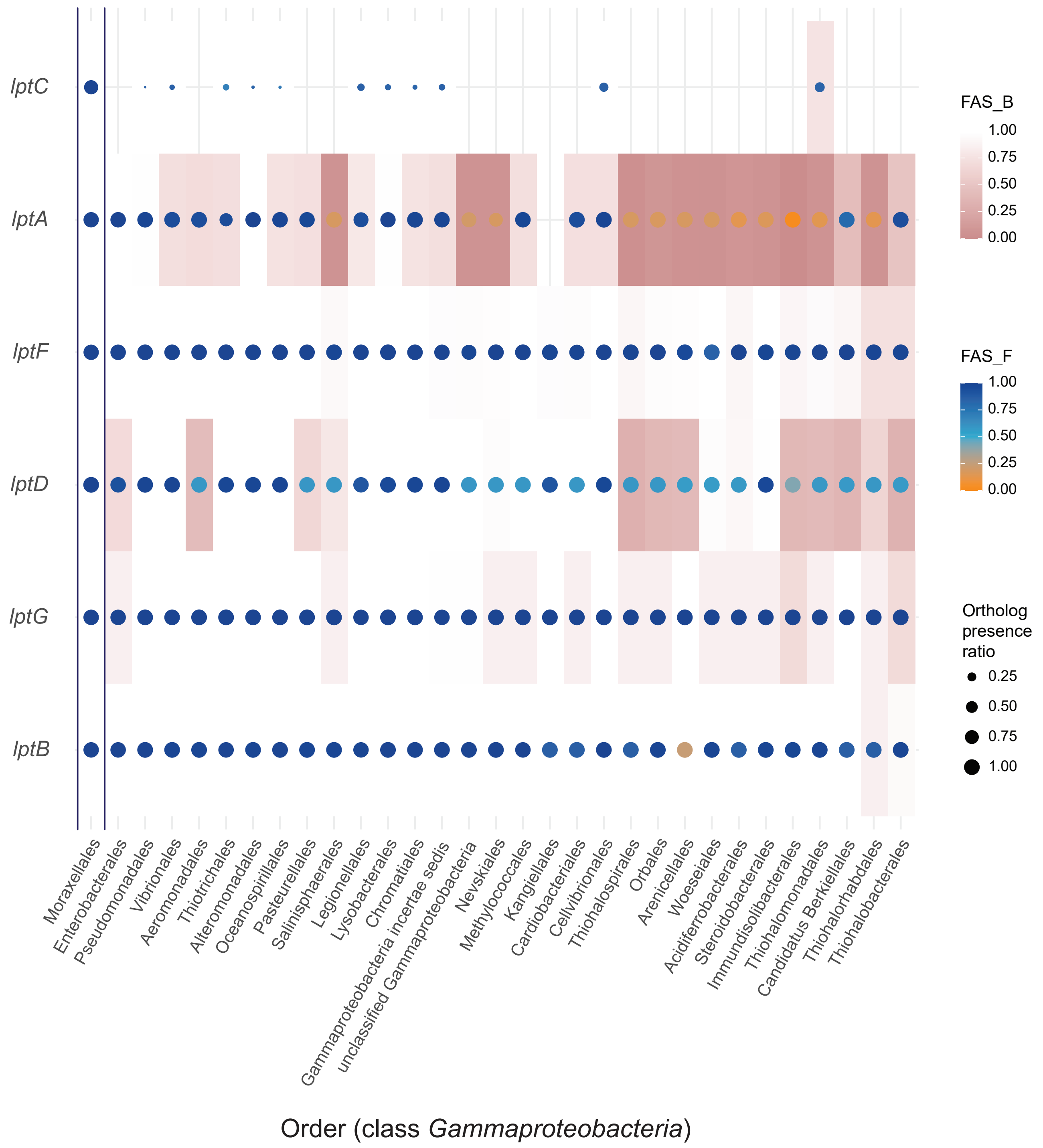

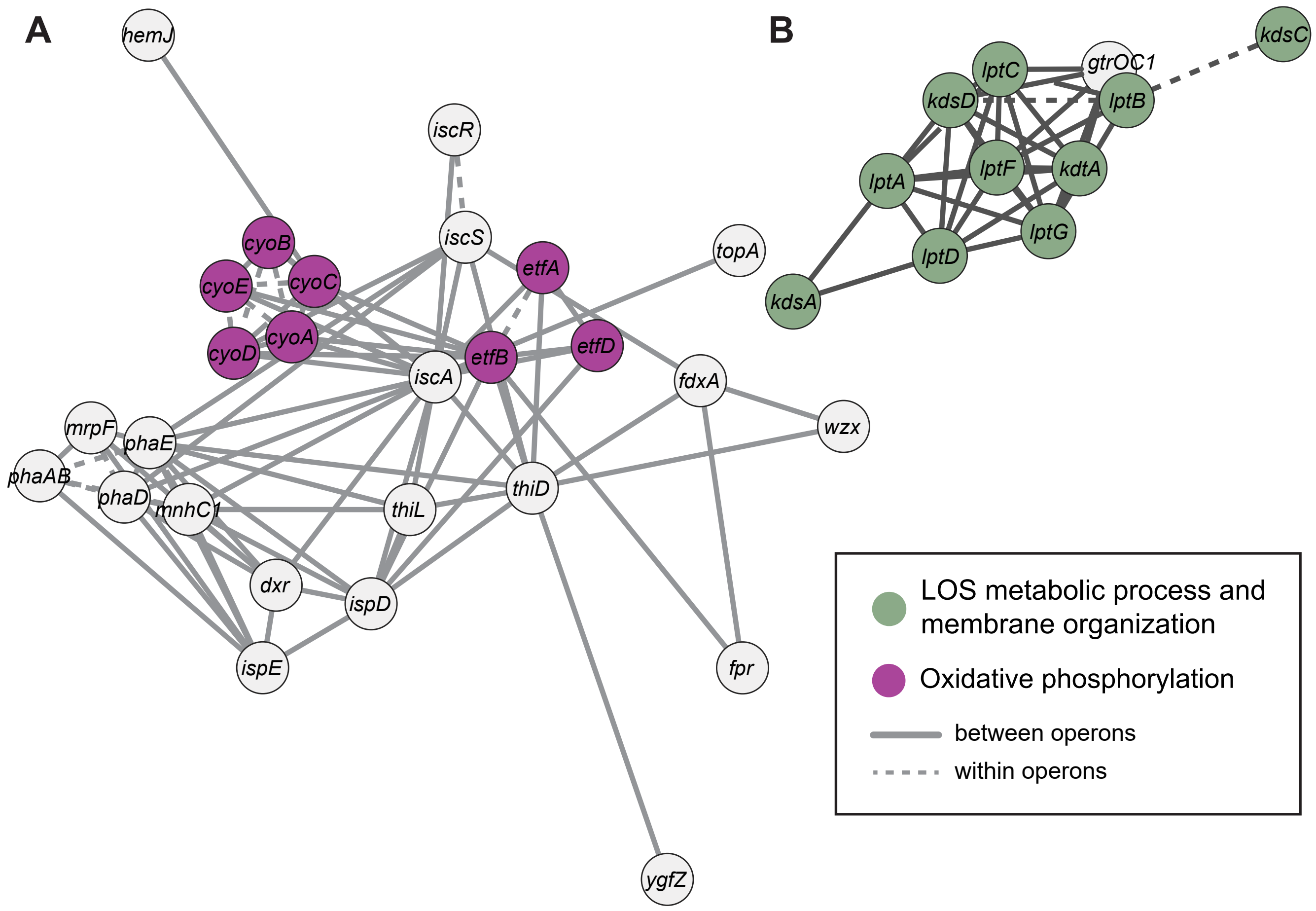

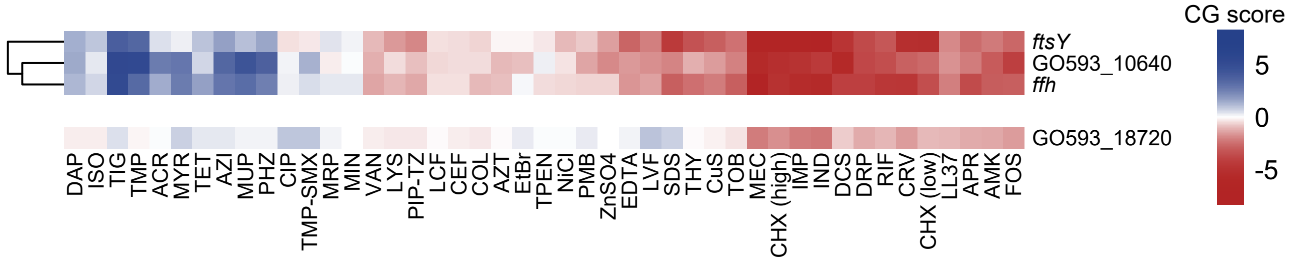

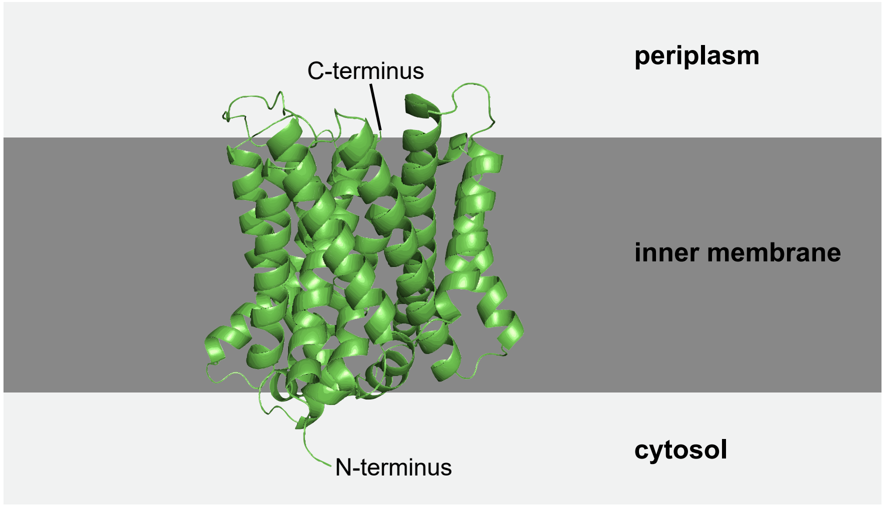

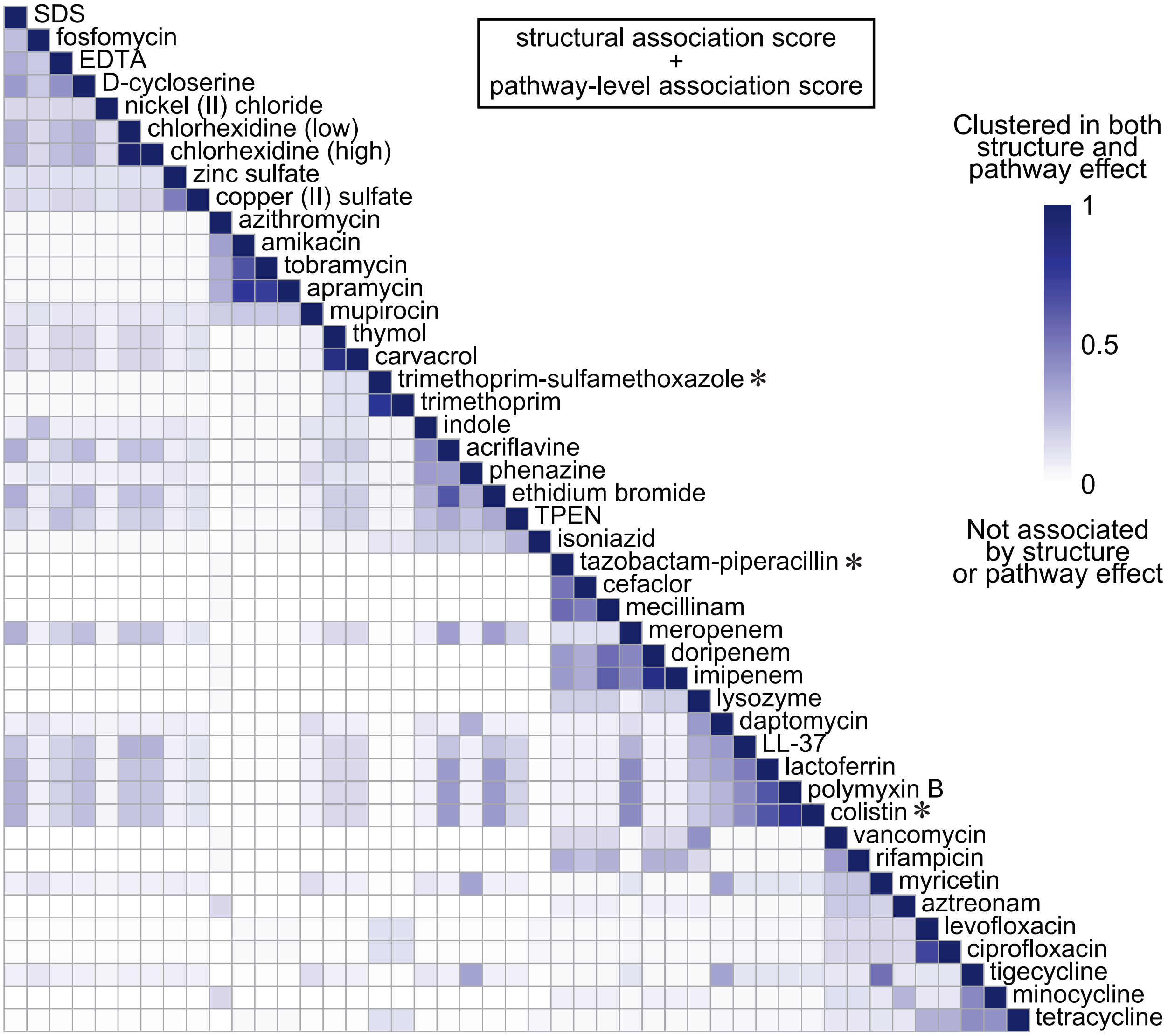

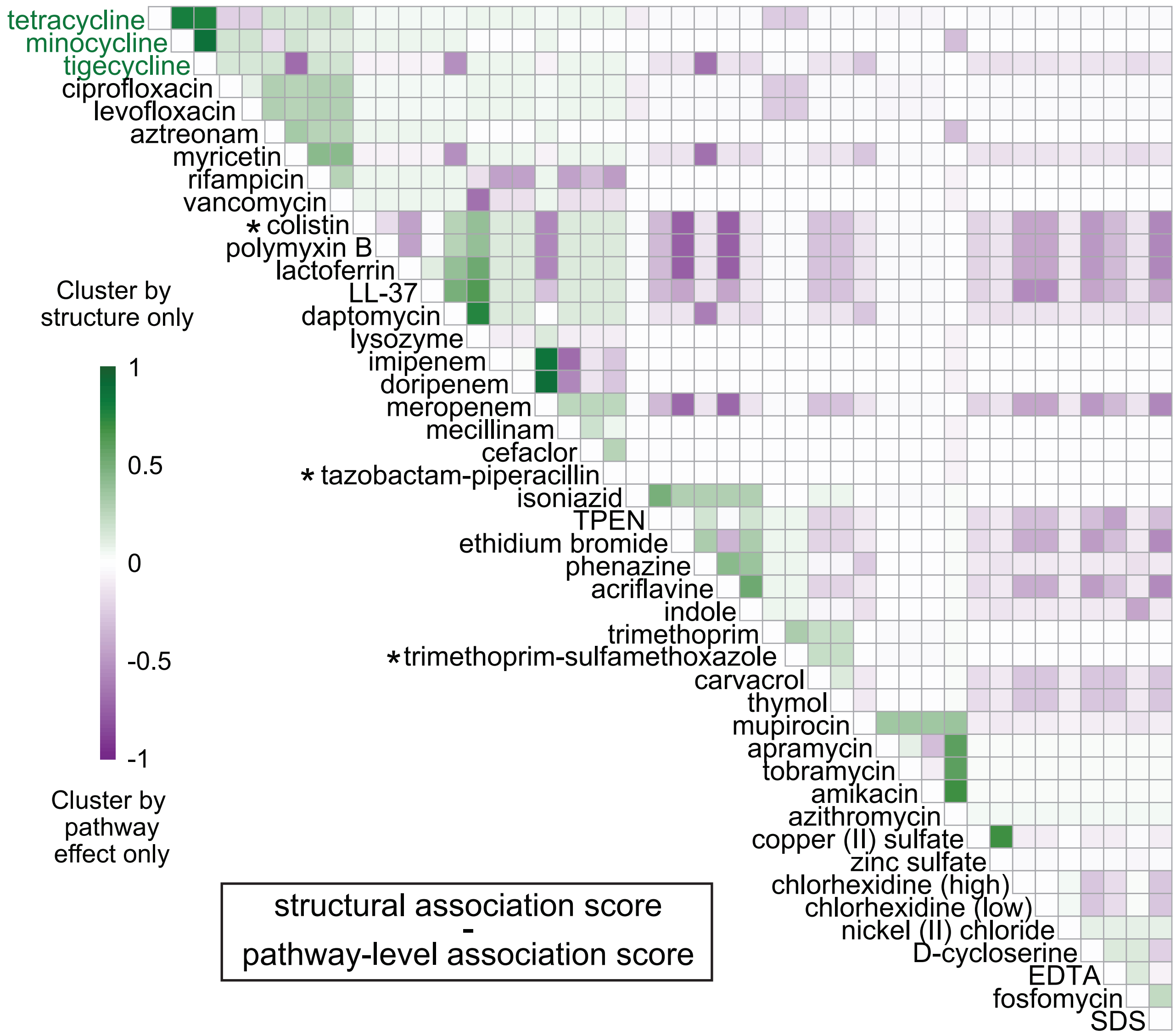

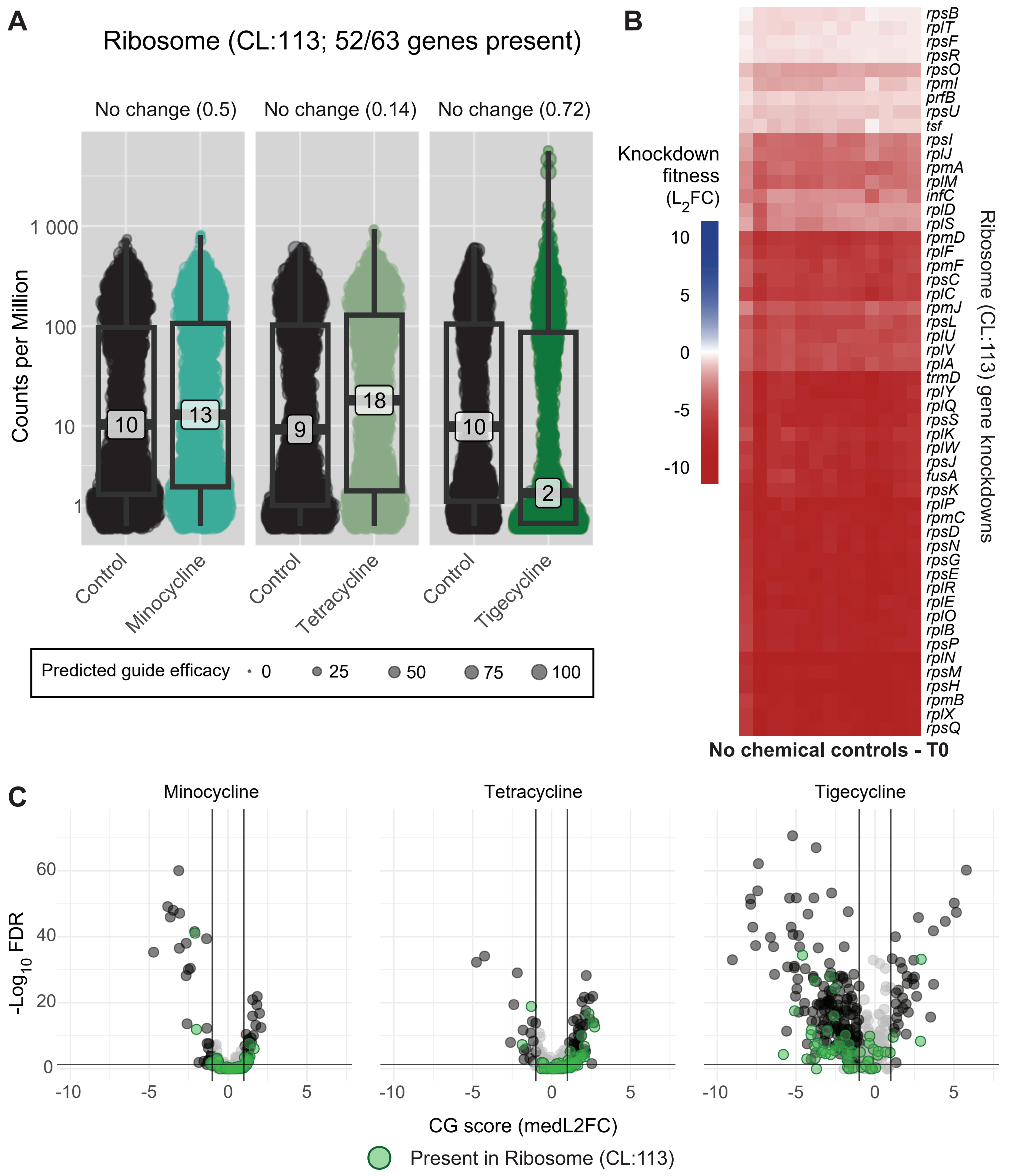

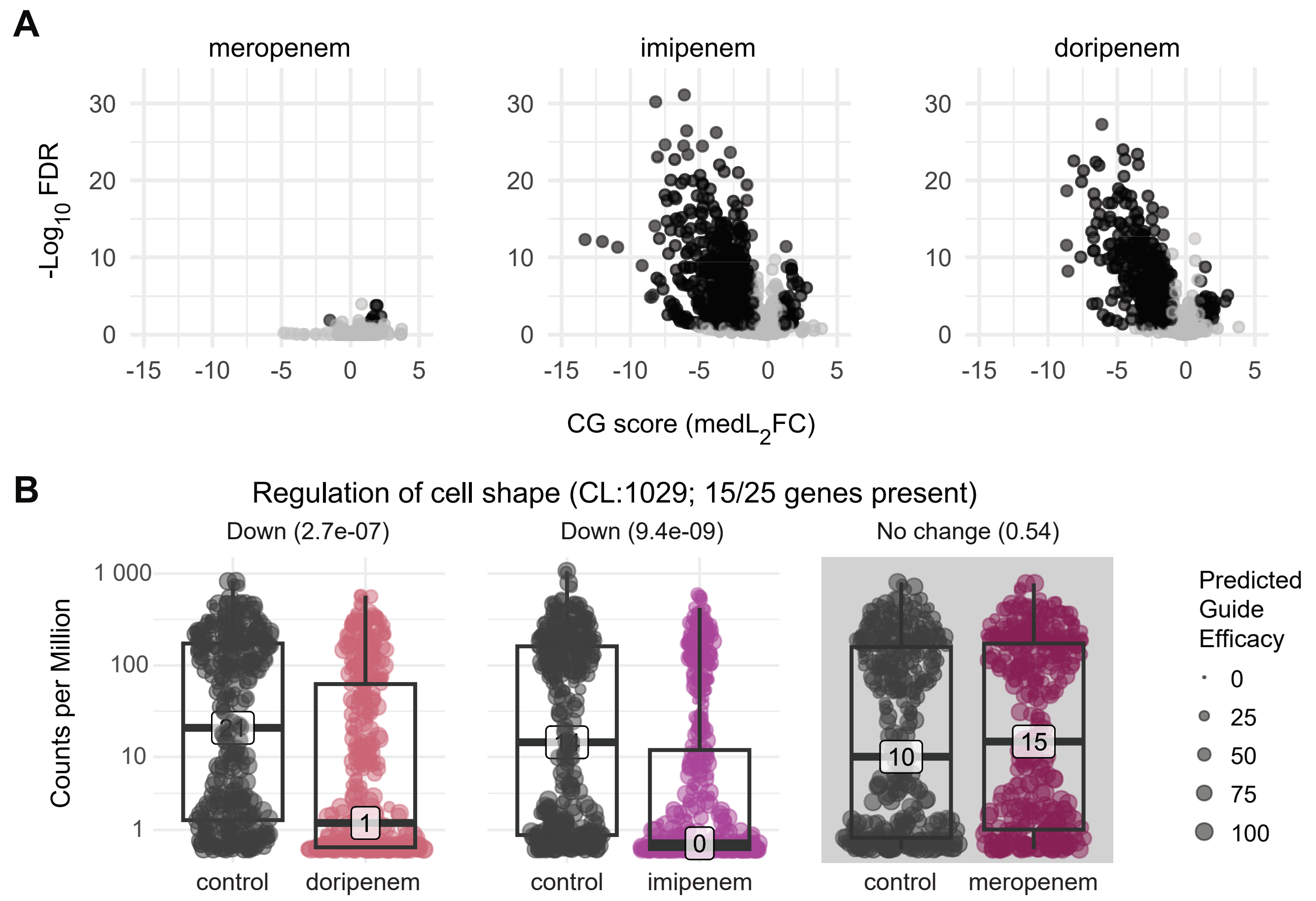

### Extended view figures

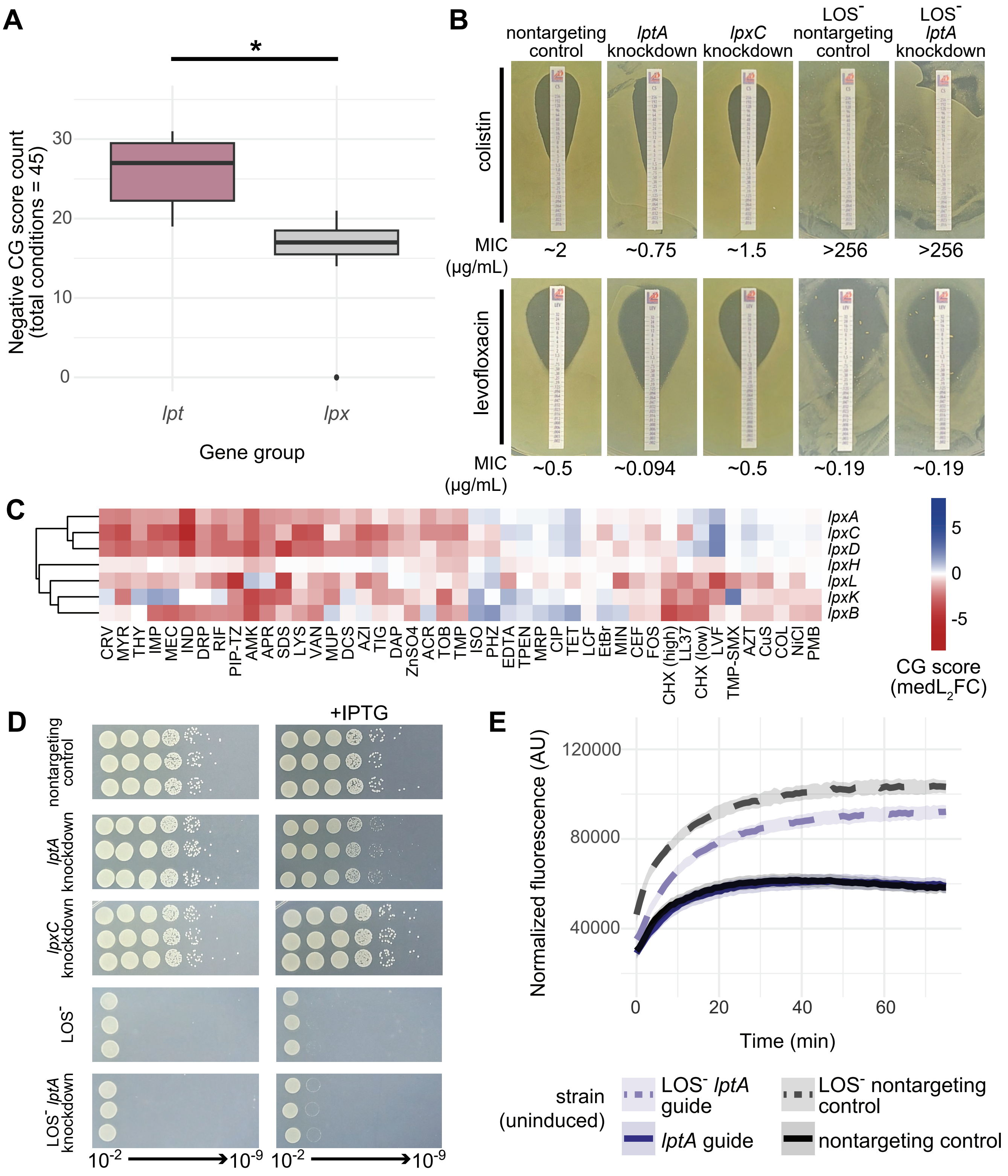

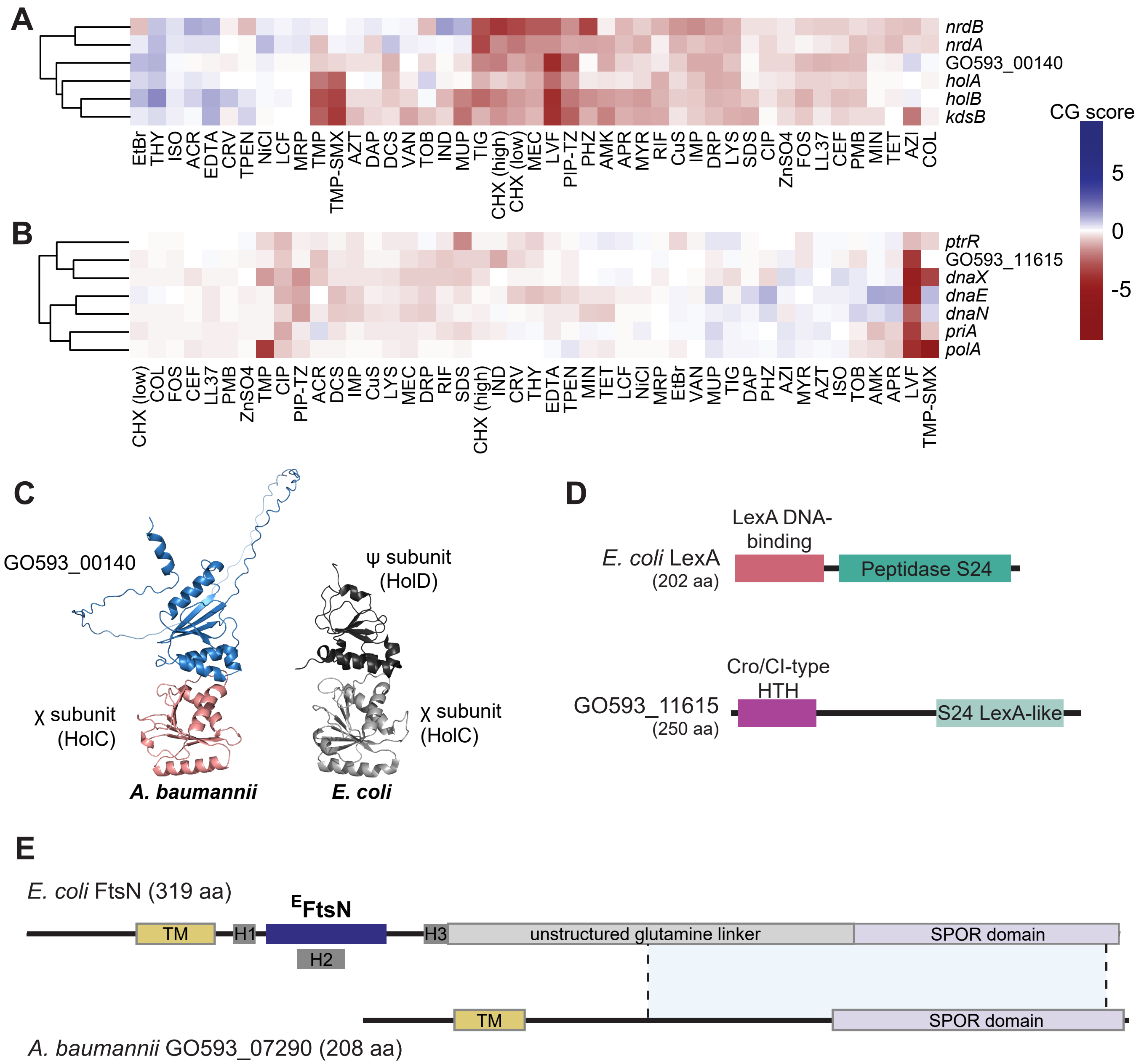

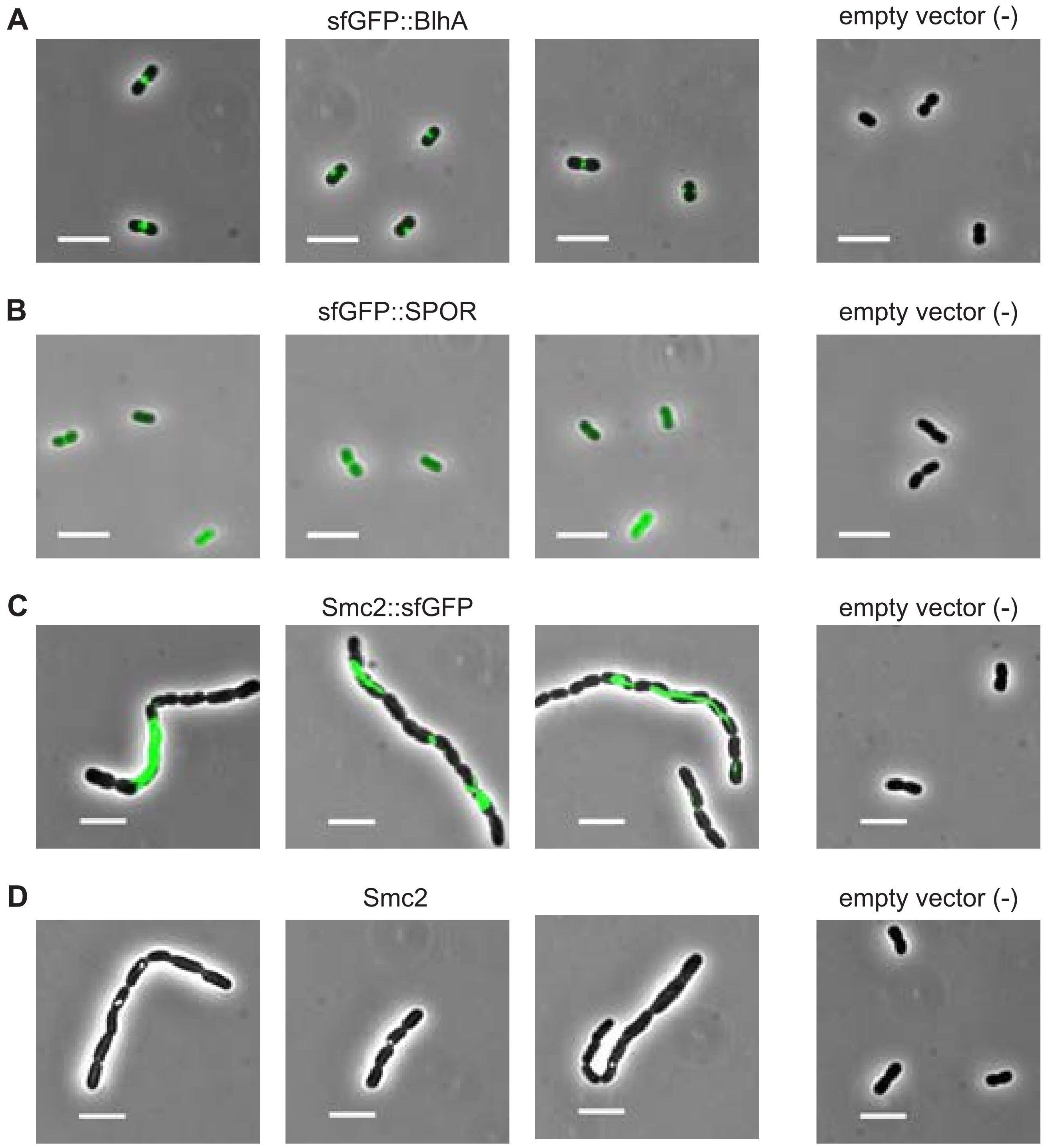

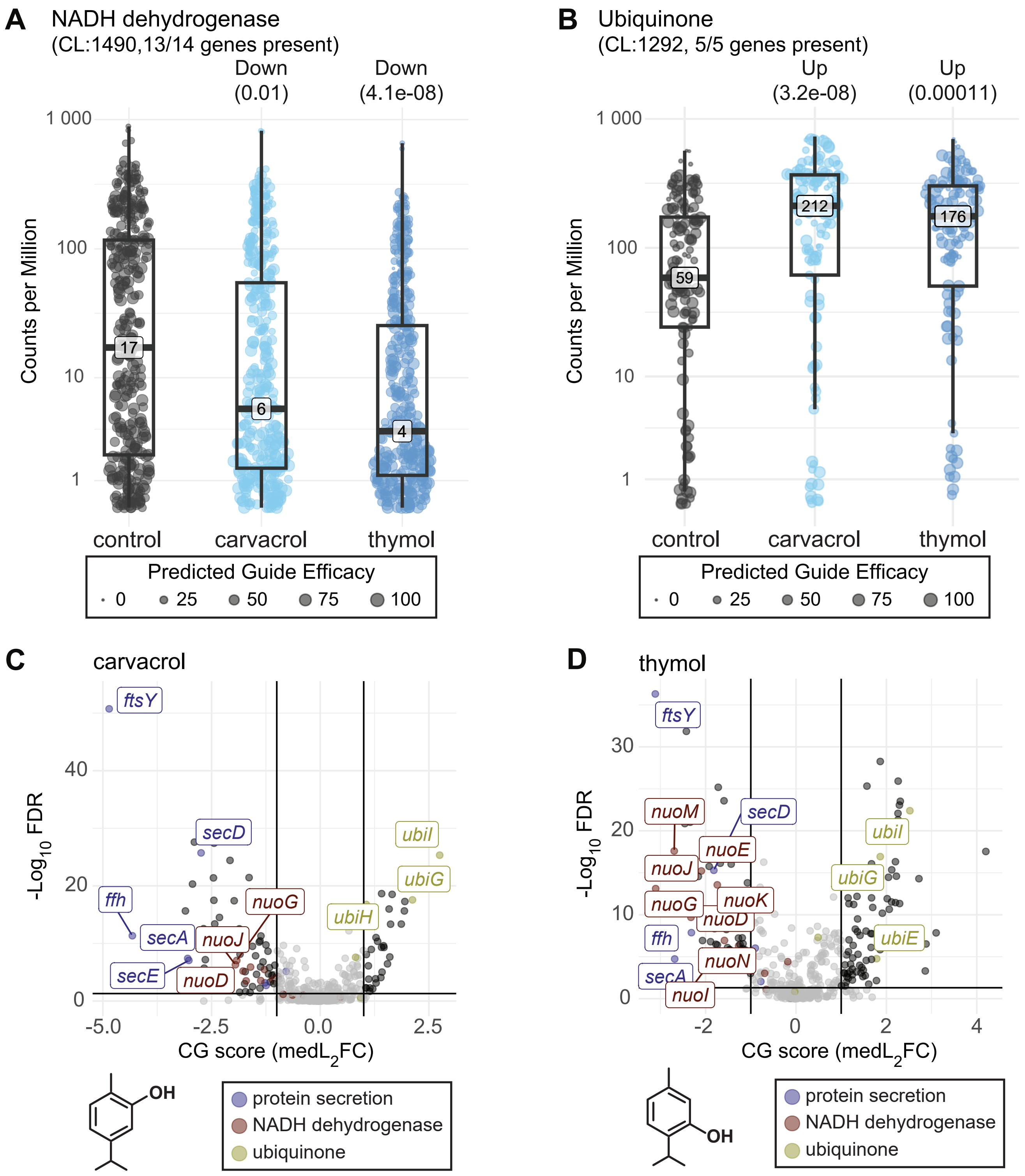

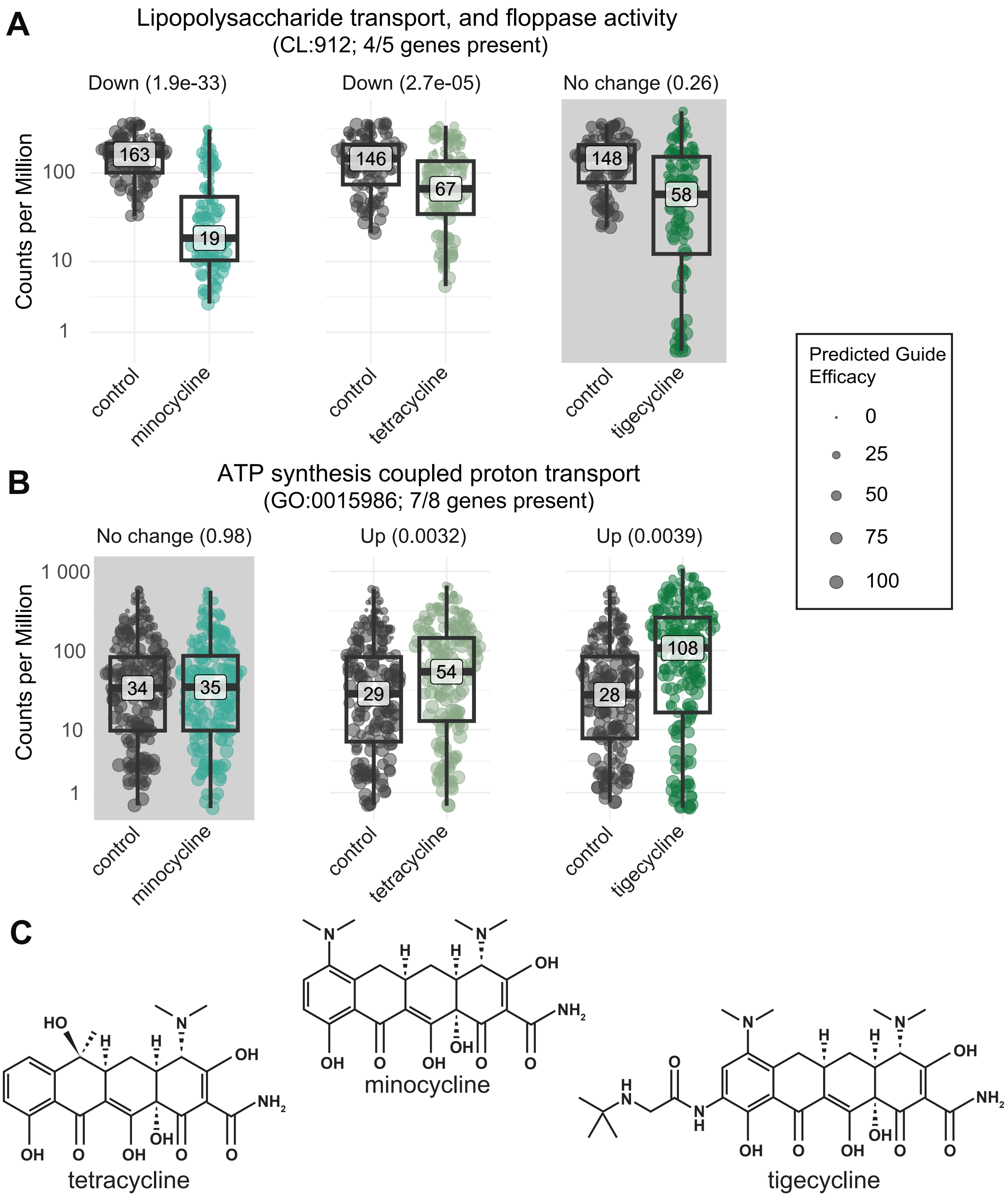
