## Supplementary material for "Chemical genomics informs antibiotic and essential gene function in *Acinetobacter baumannii*": EV+Appendix figure legends

### EXTENDED VIEW FIGURES

**Figure EV1. Growth and permeability assays for *lpt* and *lpx* knockdowns.** (A) Box plot of significant negative CG scores for *lpt* or *lpx* genes. Asterisk denotes $p < 0.05$ (Student's *t*-test). (B) Colistin and levofloxacin MIC test strip assays for *lptA* or *lpxC* knockdown, $LOS^-$, or nontargeting guide control strains. Plates supplemented with 1mM IPTG. Approximate MICs indicated below images. (C) Heatmap of CG scores of *lpx* knockdowns across conditions, showing muted phenotypes compared to *lpt* knockdowns. (D) 10 uL spots of ten-fold serial dilutions on plates with and without induction (N=3 biological replicates). The *lptA* knockdown, but not the *lpxC* knockdown, exhibits a minor growth defect upon induction. $LOS^-$ strains show major growth defects. (E) Ethidium bromide permeability assay for *lptA* knockdown and nontargeting control in 19606 or LOS- backgrounds without induction; increased fluorescence over time indicates membrane permeability. Ribbons represent standard deviation (N=4 biological replicates). Without induction, the strain containing the *lptA* guide behaves similarly to a nontargeting guide control.

**Figure EV2. Phenotypes and structural analyses of uncharacterized *A. baumannii* genes.** Heatmaps of CG scores for clusters containing (A) GO593_00140 or (B) GO593_11615. Y-axis clustering was conducted using the Ward method and Canberra distance across all library knockdowns. (C) Heterodimer structural prediction (Alphafold2) for GO593_00140 and the chi subunit of DNA Polymerase III in *A. baumannii* (left) and the solved structure for the *E. coli* psi-chi subunit heterodimer (right). GO593_00140 has a lengthy unstructured N-terminal end absent in *E. coli* HolD. (D) Domain architecture map from InterProScan analysis, comparing *E. coli* LexA to GO593_11615. Both have a DNA-binding domain and S24 peptidase/LexA domain but differ in overall protein length and specific type of DNA-binding domain. (E) Domain architectures for *E. coli* FtsN and *A. baumannii* GO593_07290. GO593_07290 lacks similarity to the short $^{E}$FtsN region required for function (47). BLAST-aligned amino acid sequences (24% identity) are depicted with dotted lines.

**Figure EV3. Microscopy of fluorescently tagged cell division proteins.** 3 representative images for each construct are shown with corresponding empty vector controls. (A) sfGFP::BlhA localizes at sites of division at the mid-cell at either existing or forming septa. (B) Fluorescently tagged SPOR-domain protein shows localization at the cell membrane and at septa for dividing cells. (C) Smc2::sfGFP expression leads to elongated cells, with fluorescently tagged proteins aggregating inconsistently within the cells. (D) Expression of Smc2 from a replicative plasmid produces elongated cells. All bars are 5 µm; GFP fluorescence/phase contrast channels merged in composite images for all except (D).

**Figure EV4. Pathways impacted by carvacrol and thymol.** Sina plots depict differential guide CPMs for (A) NADH dehydrogenase and (B) ubiquinone pathways in carvacrol and thymol. Chemical-gene interactions are described above each graph—up (positive), down (negative), or no change—with FDR values. Guides are weighted in this calculation by predicted efficacy, with perfect guides at 100. Volcano plots show relative fitness and FDRs at the gene-level in (C) carvacrol or (D) thymol. Genes in pathways of interest are highlighted, and chemical structures are depicted.

**Figure EV5. Lpt and ATP synthesis pathway effects in tetracycline-class antibiotics.** Sina plots with boxplots of differential guide CPMs for listed STRING identifiers in tetracycline-class antibiotics: (A) lipopolysaccharide (lipooligosaccharide in *A. baumannii*) transport or (B) ATP synthesis pathways. Guides are weighted by predicted efficacy, with perfect guides at 100. Chemical interactions compared to no chemical control (up, down, or no change) and FDRs are listed. Non-significant comparisons are shown with grey backgrounds. (C) Chemical structures of tetracycline, minocycline, and tigecycline.

**APPENDIX FIGURES**

**Figure S1. Replicate agreement and sample diversity as quality metrics.** (A) Correlation of library spacer counts per million (CPM) across the two biological replicates for samples containing polymyxin B at concentrations of 0.5 ug/mL (left) or 1 ug/mL (right). Count density represented by contour map; right graph shows over-depletion of guides. (B) Population diversity ($N_b$) for polymyxin B replicates. Samples with nontargeting guide complexity below the cutoff (dotted line) were excluded from analyses. (C) Heatmap showing correlations of spacer CPMs between solvent controls. All solvent control samples show $r \geq 0.97$.

**Figure S2. Chemical-gene scores and conservation in *Moraxellales*.** Dot plot depicting significant CG scores and ortholog presence ratio (fraction of isolates possessing at least one ortholog out of the total number of analyzed isolates) across representative non-*Acinetobacter Moraxellales* for library essential genes. Dots in blue represent genes with <38% presence across representative groups and significant chemical-gene interactions in >10% of screen conditions.

**Figure S3. Phyloprofile of *A. baumannii* candidate genes rare in other *Gammaproteobacteria.*** Phylogenetic profile of candidate genes showing ortholog presence ratios (fraction of isolates possessing at least one ortholog out of the total number of analyzed isolates; circle sizes) for selected *A. baumannii* genes across *Gammaproteobacteria* orders. The color encodes the median feature architecture similarity (FAS score) between the protein in 19606 and its orthologs within an order. The dot color gradient (FAS_F; blue to orange) captures architecture differences using the 19606 protein as reference. The cell color gradient (FAS_B; white to pink) captures architecture differences using the ortholog as reference. The score decreases if features in reference are missing in the respective orthologs.

**Figure S4. Phyloprofile of *lpt* genes.**
Phylogenetic profile of candidate genes showing ortholog presence ratios (fraction of isolates possessing at least one ortholog out of the total number of analyzed isolates; circle sizes) for selected *lpt* genes across *Gammaproteobacteria* orders. The color encodes the median feature architecture similarity (FAS score) between the protein in 19606 and its orthologs within an order. The dot color gradient (FAS_F; blue to orange) captures architecture differences using the 19606 protein as reference. The cell color gradient (FAS_B; white to pink) captures architecture differences using the ortholog as reference. The score decreases if features in reference are missing in the respective orthologs.

**Figure S5. Essential gene subnetworks.** Sections from the essential gene network for (A) cytochrome bo*3* oxidase (*cyo*), ion transporter (*phaAB, mnhC1, phaD, mrpF*), and related genes or (B) *lpt*-associated genes, including *gtrOC1*. Colors indicate STRING functional groups; solid or dotted lines indicate genes between or within operons, respectively.

**Figure S6. Functional phenotypes for network genes of unknown function.** (A) Heatmaps display medL2FC for the signal-recognition cluster containing *ftsY, ffh,* and GO593_10640. GO593_18720 does not cluster in the heatmap with these genes due to muted phenotypes but is connected in the network. (B) GO593_10640 is a predicted transmembrane protein similar to a transporter. Predicted structure (Alphafold) and orientation in the membrane (TMHMM) are shown.

**Figure S7. Chemical pairs similar in pathway effect and structure.** Heatmap shows the normalized sum of pathway effect and structure association scores for chemical pairs. Axes are identical. Dark blue represents chemical pairs clustered tightly in both structure or pathway effect; white represents chemicals not clustered in either structure or pathway effect. Stars represent compounds or treatments with multiple chemical structures, where one was selected for structural comparisons (see Table S4).

**Figure S8. Chemical pairs opposed in pathway effect and structure.** Heatmap shows the normalized differences between pathway effect and structure association scores for chemical pairs. Axes are identical. Green represents chemical pairs clustered tightly by structure but not pathway effect; purple represents chemicals clustered by pathway effect but not by structure. Stars represent compounds or treatments with multiple chemical structures, where one was selected for structural comparisons (see Table S4).

**Figure S9. Target pathway effects for tetracycline-class antibiotics.** (A) Sina plots depict differential guide CPMs for ribosome genes (STRING identifier CL:113) in minocycline, tetracycline, or tigecycline compared to no chemical control. Chemical-gene interactions are described above each graph—up (positive), down (negative), or no change—with FDR values. Guides are weighted in this calculation by predicted efficacy, with perfect guides at 100. (B) Heatmap shows relative fitness scores(median $\log_2$ fold changes) of ribosomal gene knockdowns of the mock treatment control compared to T0. Ribosome knockdowns all have substantial loss of fitness without additional chemical treatment. (C) Volcano plots of library gene CG scores (median $\log_2$ fold change compared to mock treatment) in minocycline, tetracycline, and tigecycline. Genes in the ribosome group are colored green. Lines depict cutoffs for significance (|median $\log_2$ fold change| ≥1, Stouffer's $p < 0.05$).

**Figure S10. Ineffective chemical concentrations lack significant pathway effects.** (A) Volcano plots of genes in meropenem, imipenem, or doripenem compared to no drug control. Genes with significant CG scores are shown in black. Meropenem has very few significant CG scores, suggesting insufficient chemical dosage. (B) Sina plots with boxplots of differential guide CPMs for target pathway in doripenem, imipenem, or meropenem. Chemical interactions compared to no chemical control (up, down, or no change) and FDRs are listed. Non-significant comparison for meropenem is shown with grey background. Doripenem and imipenem are significantly impacted.
